# FRIENDLY (FMT) is an RNA binding protein associated with cytosolic ribosomes at the mitochondrial surface

**DOI:** 10.1101/2022.01.27.478018

**Authors:** Mickaele Hemono, Thalia Salinas-Giegé, Jeanne Roignant, Audrey Vingadassalon, Philippe Hammann, Elodie Ubrig, Patryk Ngondo, Anne-Marie Duchêne

## Abstract

The spatial organization of protein synthesis in the eukaryotic cell is essential for maintaining the integrity of the proteome and the functioning of the cell. Translation on free polysomes or on ribosomes associated with the endoplasmic reticulum has been studied for a long time. More recent data have revealed selective translation of mRNAs in other compartments, in particular at the surface of mitochondria. Although these processes have been described in many organisms, in particular in plants, the mRNA targeting and localized translation mechanisms remain poorly understood.

Here, the *Arabidopsis thaliana* Friendly (FMT) protein is shown to be a cytosolic RNA binding protein that associates with cytosolic ribosomes at the surface of mitochondria. Its knockdown delays seedling development and causes mitochondrial clustering. The mutation also disrupts the mitochondrial proteome and the localization of nuclear transcripts on the surface of mitochondria. These data indicate that FMT participates in the localization of mRNAs and their translation at the surface of mitochondria.

## INTRODUCTION

Mitochondria are vital organelles for the eukaryotic cell, through their role in ATP synthesis and other metabolic pathways. Their biogenesis is based on the expression of the mitochondrial genome but also relies on a massive import of nuclear-encoded proteins, which are translated in the cytosol. It has been recently shown that protein import can occur post-translationally and co-translationally. In accordance, cytosolic mRNAs and ribosomes were found associated with the mitochondrial surface in numerous organisms.

One third to one half of mRNAs coding for nuclear-encoded mitochondrial proteins were found at the surface of mitochondria in plants (Vincent et al., 2017), yeast (Saint-Georges et al., 2008; Williams et al., 2014), or mammals (Fazal et al., 2019). The mechanisms and controls governing mRNAs targeting to the mitochondrial surface are poorly understood. Both nucleotide motifs in the transcript and sequence signals in the newly synthesized protein can be important for mitochondrial surface localization of mRNAs (Lesnik et al., 2015; Lashkevich and Dmitriev, 2021).

Cytosolic ribosomes near the mitochondrial outer membrane were first observed in yeast by Butow’s team in the ‘70s (Kellems et al., 1974), and was confirmed more recently by electron cryo-tomography (Gold et al., 2017). The precise mechanism of cytosolic ribosomes recruitment at the surface of mitochondria is not clear, but the Nascent polypeptide Associated Complex (NAC) (Funfschilling and Rospert, 1999) and the mitochondrial outer membrane protein OM14 (Lesnik et al., 2014) were shown involved in yeast, as well as the Translocase of the Outer Membrane (TOM) complex through the interaction with the targeting sequence of the newly synthesized protein (Gold et al., 2017).

Some RNA-binding proteins (RBPs) have been found involved in mRNA localization and translation at the mitochondrial surface, such as PUF3 in yeast ((Saint-Georges et al., 2008) or Larp in Drosophila (Zhang et al., 2016). The human CLUH was also shown to interact with mRNAs coding for mitochondrial proteins (Gao et al., 2014), and its drosophila orthologue, Clueless, was able to bind ribosomal components at the surface of mitochondria (Sen and Cox, 2016). More recently, the CLUH interactome revealed the proximity of CLUH with mitochondrial proteins and their mRNAs during cytosolic translation (Hemono et al., 2022). CLUH and Clueless belong to the CLU family, which is widespread in all eukaryotes. CLU stands for “CLUstered mitochondria” because knock-down of CLU proteins induces clustering of mitochondria in all studied organisms (El Zawily et al., 2014).

In plants, the specificities of cytosolic ribosomes at the surface of mitochondria are not known. Moreover, no RBP involved in mRNA localization and localized translation has been identified. To characterize ribosomes at the surface of the plant mitochondria, we have purified cytosolic ribosomes from an *Arabidopsis thaliana* mitochondrial extract. We have found that Friendly (FMT), the *A. thaliana* CLUH orthologue, was specifically enriched in this ribosomal fraction. Using reverse co-immunoprecipitation and split GFP approaches, we have confirmed the association of FMT with the cytosolic ribosome and with the mitochondrial surface. We have also established that FMT is an RBP. We have observed that *fmt* knock-down caused a reduced growth of seedlings and a clustering of mitochondria. At the level of mitochondria, the *fmt* KO mutation was found to affect the mitochondrial proteome and the association of cytosolic mRNAs with mitochondria, suggesting a role of FMT in mRNA localization at the mitochondrial surface.

## RESULTS

### FMT copurifies with cytosolic ribosomes isolated from mitochondrial extracts

To characterize particular components of the cytosolic ribosome at the mitochondrial surface, an *Arabidopsis thaliana* (*At*) line expressing an epitope-tagged cytosolic ribosomal protein, FLAG-RPL18 (AT3G05590), was used (Zanetti et al., 2005). Total and mitochondrial proteins extracts were prepared from FLAG-RPL18 and wild-type (Col0) inflorescences. Inflorescences were chosen because they are mitochondria-enriched tissues (Welchen et al., 2014). The quality of mitochondrial extracts was verified with western blots (**Figure S1**). Proteins were then immunoprecipitated with anti-FLAG beads (IP), identified by LC-MS/MS analyses, and spectral count label-free quantifications were performed. First, the IP with mitochondrial extracts were compared, that is IP with Flag-<u>R</u>PL18 <u>m</u>itochondria (Rm-IP) and with <u>C</u>ol0 <u>m</u>itochondria (Cm-IP) (**Figure 1A**), and 185 proteins were found enriched in Rm-IP (with cuts-off of Fold Change (FC) above 2 and adjusted p-value (adj-p) below 0.1). One hundred and eleven out of the 185 proteins were from the cytosolic ribosome, confirming the presence of cytosolic ribosomes in the mitochondrial extract. IP from <u>m</u>itochondrial and <u>t</u>otal extracts from the FLAG-<u>R</u>PL18 line were also compared (Rm-IP versus Rt-IP, **Figure 1B**), and 96 proteins were found enriched in Rm-IP.

**Figure 1:**
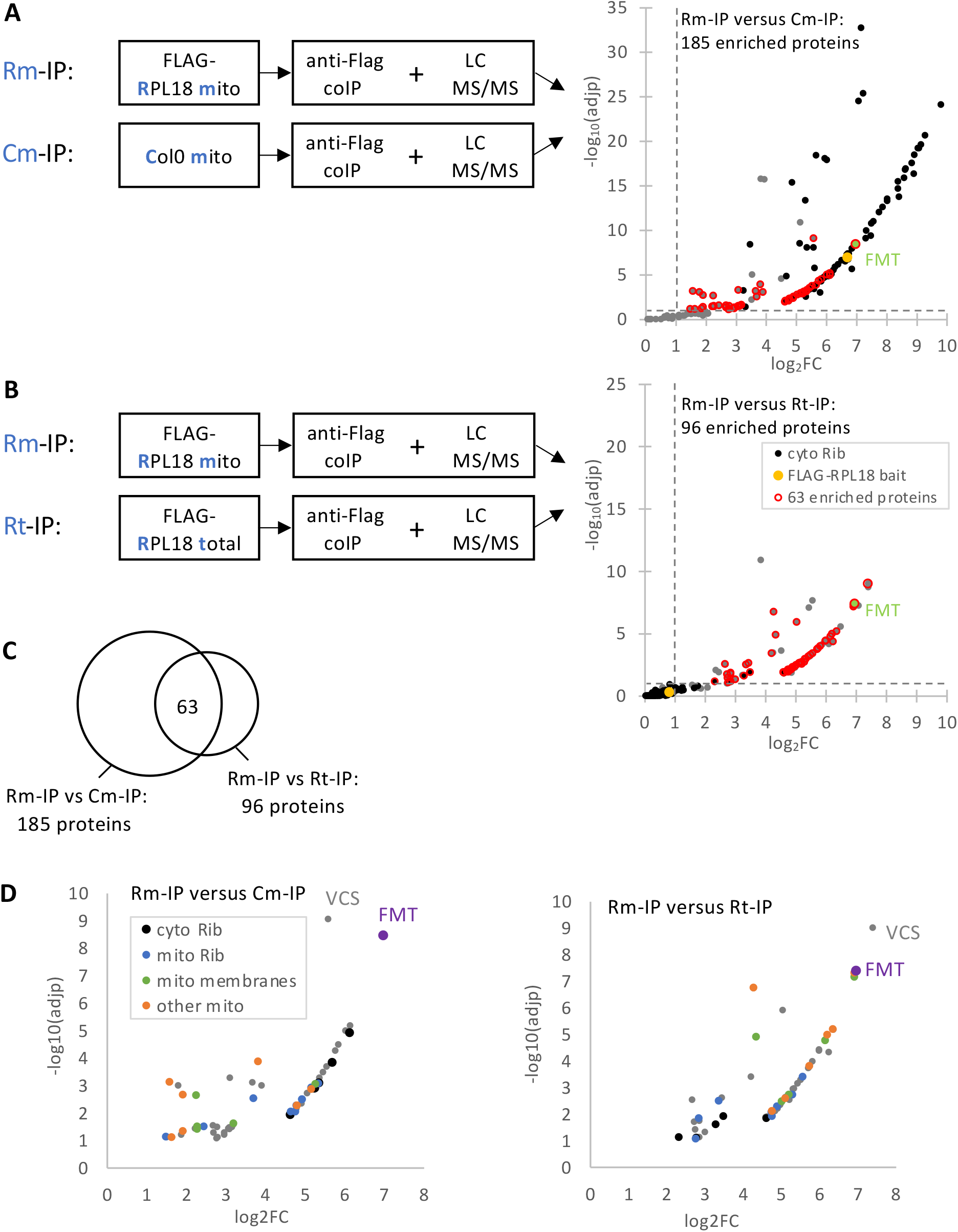
Identification of proteins co-purifying with FLAG-RPL18 at the surface of mitochondria. **A**-FLAG-RPL18 mitochondrial coIP compared to Col0 mitochondrial one (Rm-IP versus Cm-IP). Immunoprecipitated proteins were identified by LC-MS/MS. Statistical analyses based on specific spectral counts identified 185 proteins enriched in Rm-IP compared to Cm-IP with cuts-off of Fold Change (FC) above 2 and adjusted p-value (adj-p) below 0.1 **B**-FLAG-RPL18 mitochondrial coIP compared to FLAG-RPL18 total one (Rm-IP versus Rt-IP). With the same cuts-off as in A, 96 proteins were enriched in Rm-IP compared to Rt-IP. In A and B, the yellow spots correspond to the RPL18 bait. Proteins from cytosolic ribosomes (Salih et al., 2020) are in black. The 63 enriched proteins (see C) are with a red circle and FMT is in green. **C**-Sixty-three enriched proteins are shared in these two comparisons, Rm-IP versus Cm-IP (A), and Rm-IP versus Rt-IP (B). **D**-Graphs in A and B are enlarged for the 63 proteins. Five out of the 63 proteins are from the cytosolic ribosome (black). Among mitochondrial proteins, 5 are components of the mitochondrial membranes (green), 10 are from the mitochondrial ribosome (blue). Mitochondrial ribosomes are known to be associated with membranes (Tomal et al., 2019); Waltz and Giege, 2020). The other mitochondrial proteins are in orange.

Sixty-three proteins appeared shared between the 2 comparisons, that is enriched in Rm-IP compared to both Cm-IP and Rt-IP (**Figure 1C-D Table S1**). Five out of these 63 proteins have been identified as components of the cytosolic ribosome (Salih et al., 2020). Twenty-three others were annotated as mitochondrial according to SUBA4 (Hooper et al., 2017). No proteins from TOM complex were found, and only one protein from the Mitochondrial Outer Membrane (MOM), the voltage-dependent anion channel VDAC2, was identified. This contrasts with results in yeast where interaction between cytosolic ribosomes and TOM complex has been found (Gold et al., 2017).

The most enriched proteins among the 63 are 2 cytosolic proteins, VCS (AT3G13300) and FMT (AT3G52140). VCS forms an mRNA decapping complex with DCP1 and DCP2 in processing bodies (P-bodies) (Xu and Chua, 2011). However, neither DCP1 (AT1G08370), nor DCP2 (AT5g13570) could be identified in any of the above IPs. FMT, a member of the CLUSTERED MITOCHONDRIA (CLU) family, is required for the correct distribution of mitochondria within the cell (El Zawily et al., 2014). Because of its orthologues’ properties and its role in plant mitochondria distribution (El Zawily et al., 2014), FMT was further studied.

### FMT is an RBP that strongly interacts with the ribosome at the surface of mitochondria

To explore FMT functions, it was essential to establish its localization in the cell. Two translational fusions of FMT with GFP were constructed, one with GFP in N-terminal (GFP-FMT), the other with GFP in C-terminal (FMT-GFP). *At* lines stably expressing the 2 FMT fusions were obtained. Both FMT fusions gave a cytosolic location, often with a punctuate distribution (**Figure 2A**). When transiently expressed in *Nicotiana benthamiana* (*Nb*), the FMT constructs gave similar cytosolic signals, which were sometimes found in the vicinity of mitochondria (**Figure 2B**). Such localizations have been previously observed by others (El Zawily et al., 2014) (Ma et al., 2021; Ayabe et al., 2021), and could be linked to different aspects of FMT function.

**Figure 2:**
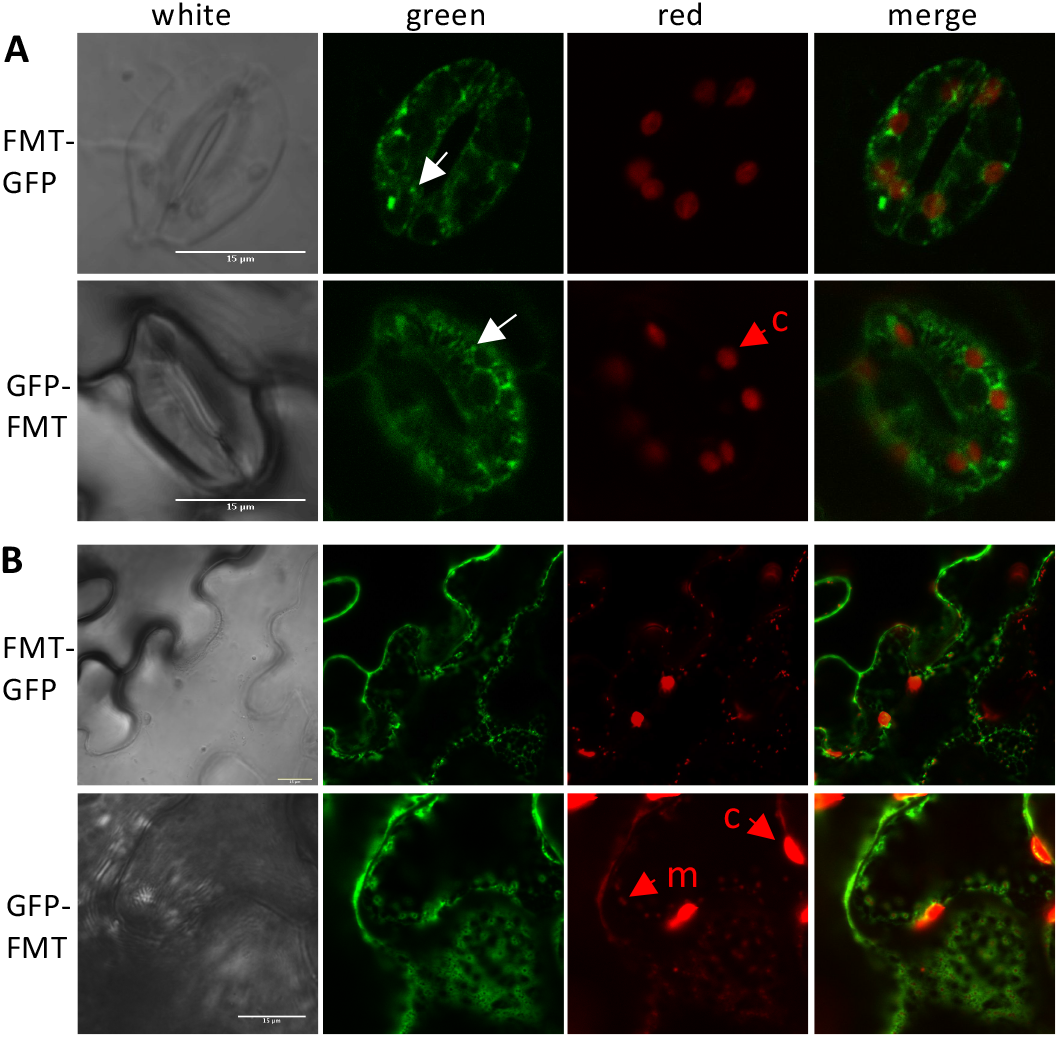
FMT localization in plant cells. **A -** *A. thaliana* 18-day-old seedlings stably expressing FMT-GFP or GFP-FMT The GFP signal showed a diffuse cytosolic localization and also a punctuate distribution (white arrows) **B -** *N. benthamiana* 6-week-old leaves transiently expressing FMT-GFP or GFP-FMT, and the mitochondrial pSU9-RFP Leaves were examined by confocal microscopy. White: bright light; green: GFP fluorescence; red: chloroplast autofluorescence, and mitochondrial pSU9-RFP fluorescence in B; merge, red and green channels. c, chloroplast; m, mitochondrium. Scale bars: 15 m

To determine FMT interactants, mitochondrial proteins extracts were prepared from inflorescences of *At* lines expressing GFP-FMT or FMT-GFP, and Col0. Proteins were immunoprecipitated with anti-GFP beads and identified by mass spectrometry. The IPs were then compared, that is GFP-FMT with Col0, and FMT-GFP with Col0 (**Figure 3 A-B**). Respectively, 73 and 28 proteins were found enriched in GFP-FMT and FMT-GFP compared to Col0, with cuts-off of Fold Change above 2 and adjusted p-value below 0.1. In total, 75 proteins were co-purified with at least one of the 2 FMT constructs (**Figure 3 C-D**; **Table S2**). Thirty-five of the 75 were cytosolic ribosomal proteins, and 25 others were mitochondrial. CoIPs with anti-GFP beads were also performed with mitochondria from seedlings of the GFP-FMT line and Col0. The co-purification of FMT and cytosolic ribosomal proteins was again obtained (**Figure S2, Table S3**). All these reverse coIPs confirmed that FMT co-purified with cytosolic ribosomes in mitochondrial fractions.

**Figure 3:**
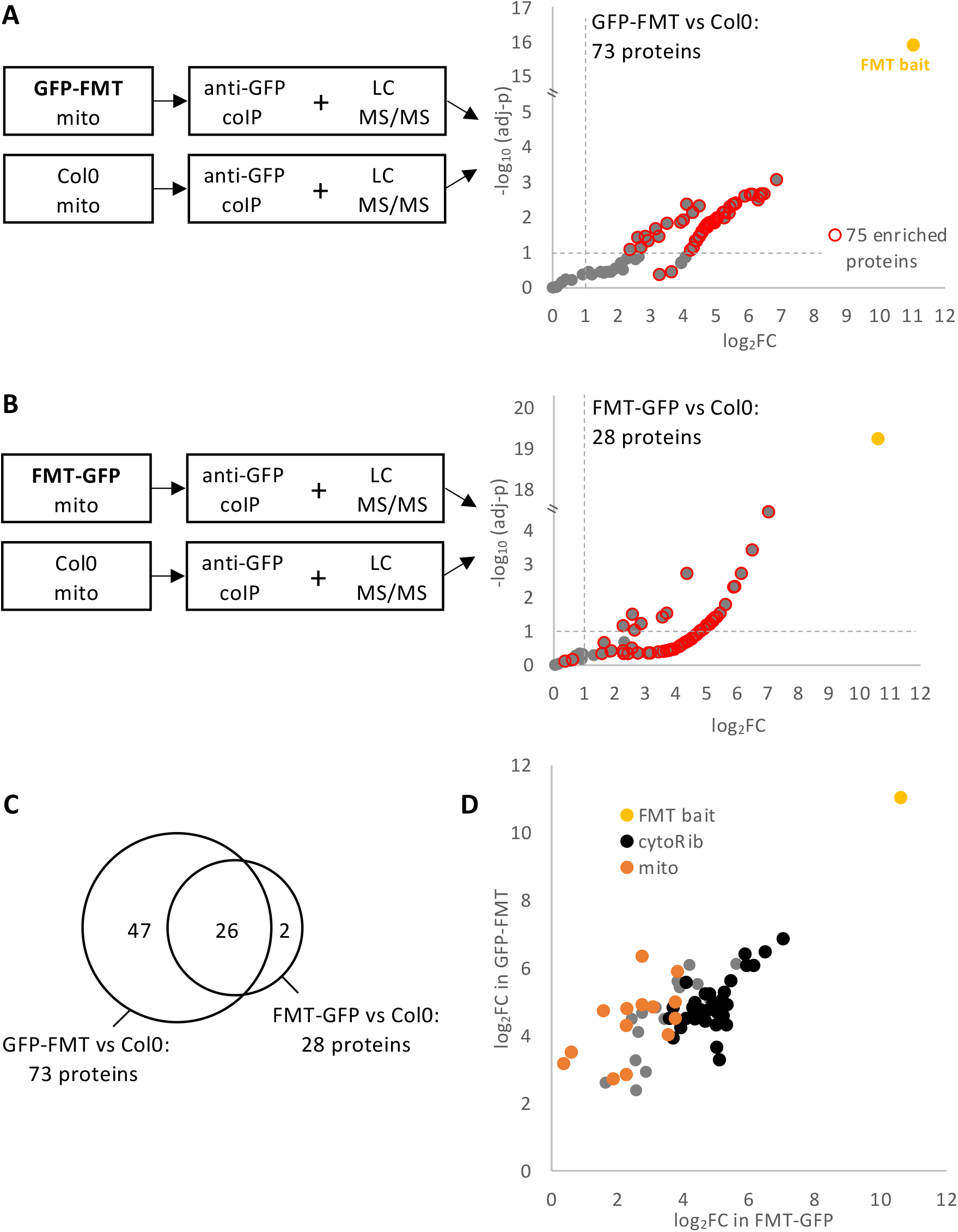
Identification of proteins co-purifying with FMT at the surface of inflorescence mitochondria. **A**-GFP-FMT mitochondrial coIP compared to Col0 mitochondrial coIP (GFP-FMT versus Col0) Immunoprecipitated proteins were identified by LC-MS/MS. Statistical analyses based on specific spectral counts identified 73 proteins enriched in FMT-GFP compared to Col0 (cuts-off FC> 2 and adj-p<0.1). **B**-FMT-GFP mitochondrial coIP compared to Col0 mitochondrial coIP (FMT-GFP versus Col0) With the same cuts-off as in A, 28 proteins were enriched in GFP-FMT compared to Col0. **C**-In total, 75 proteins were enriched in at least one FMT/GFP IP. They are shown with a red circle in A and B. **D**-The enrichment of the 75 proteins in FMT coIPs. x- and y-axes correspond respectively to log_2_FC in FMT-GFP and GFP-FMT coIPs. The FMT bait is in yellow. Proteins from cytosolic ribosomes are in black, mitochondrial proteins are in orange.

Because FMT is also cytosolic (**Figure 2**), coIPs were also performed with total extracts from inflorescences and seedlings of GFP-FMT and FMT-GFP lines, and of Col0. Only one ribosomal protein and one other protein (a mitochondrial PDH subunit encoded by AT3G13930) were found enriched in FMT coIPs. (**Figure S3 A, B**). Seedlings total extracts were also crosslinked with formaldehyde before immunoprecipitation to withstand the isolation procedure. In these conditions, 119 proteins were found enriched in FMT coIPs, and 81 were ribosomal proteins (**Figure S3C**, **Table S4**).

Altogether, these coIPs revealed that FMT poorly or partially interacted with cytosolic ribosomes in a total extract. By contrast, FMT was found to strongly interact with cytosolic ribosomes in the mitochondrial fractions.

The split-GFP approach (Romei and Boxer, 2019) was used to confirm the proximity of FMT with the ribosome and with the mitochondrial membrane. Translational fusions with either GFP-β11 strand or GFP-β1-10 strands were transitory co-expressed in *N. benthamiana* (**Figure 4, Figure S4**). The co-expression of β1-10-TOM5 and β11-FMT on the one hand, and of β1-10-RPL18 and β11-FMT on the other hand, resulted in fluorescence, showing that these proteins pairs allow the complementation of the 2 GFP fragments in *N. benthamiana.* The split-GFP approach thus confirmed the proximity of FMT with the ribosome and with the mitochondrial surface.

**Figure 4:**
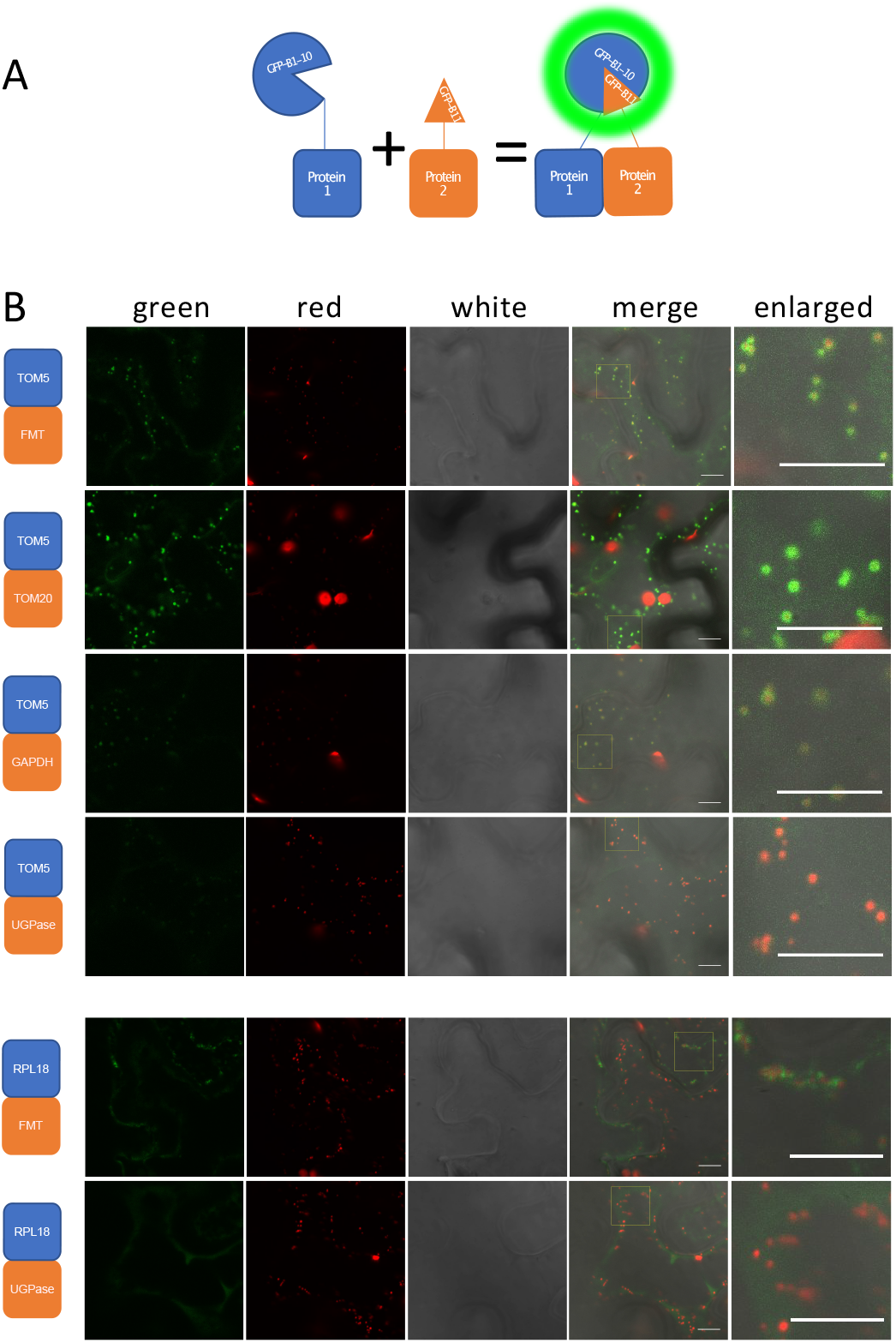
Split-GFP experiments. **A**-Schematic representation of Split-GFP experiment. The proteins of interest were respectively fused with GFP-β1-10 strands (Protein 1, in blue) or GFP-β11 strand (Protein 2, in orange). *N. benthamiana* leaves were co-transformed with constructs encoding β1-10 and β11 fusions and with the mitochondrial pSU9-RFP construct. The co-localization of the two fusion proteins allowed the reconstitution of a functional GFP. **B**-Confocal imaging. For interactions with mitochondria, the MOM TOM5 protein was fused with β1-10. The co-expression with FMT fused to β11 gave a signal around mitochondria. The TOM5-TOM20 and TOM5-GAPDH pairs served as positive controls. TOM5 and TOM20 are 2 components of the TOM complex in MOM. The cytosolic GAPDH enzyme is found at the surface of mitochondria (Giege et al., 2003), so the TOM5-GAPDH pair is a positive control for the interaction of a cytosolic protein with MOM. By contrast TOM5-UGPase served as a negative control, UGPase being only cytosolic. For interactions with the ribosome, RPL18 was fused with β1-10. The co-expression with FMT fused to β11 gave a signal often associated with mitochondria. By contrast the RPL18-UGPase pair gave a weak and diffuse signal in the cytosol. Alone, none of these constructs gave fluorescence (Figure S4). Green: GFP fluorescence; red: chloroplast autofluorescence and mitochondrial pSU9-RFP fluorescence; white: Bright light. All the scale bars correspond to 10 m.

To determine if FMT was an RBP, an oligo(dT) capture experiment was performed. Leaves from *At* GFP-FMT line were first irradiated with UV to induce covalent bonds between RNAs and interacting proteins. Then, the leaves extracts were used for the capture of poly(A) RNAs and covalently linked proteins using oligo(dT) magnetic beads. FMT was detected in the eluate, confirming its RNA binding property (**Figure 5**). The Drosophila and mammalian FMT orthologues were shown as RBPs (Sen and Cox, 2016; Gao et al., 2014).

**Figure 5:**
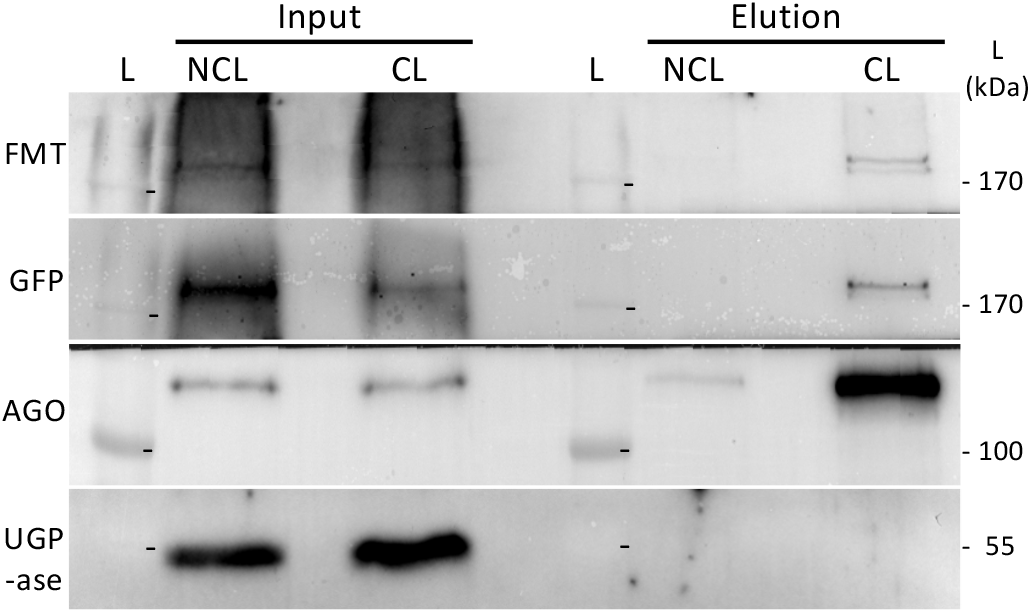
FMT is an RNA-binding protein. Leaves from *At* GFP-FMT line were UV-irradiated to crosslink RNA and proteins (CL fractions). The crosslinked mRNA-protein complexes were pulled down by oligo-d(T)_25_ beads. A similar experiment was performed with non-crosslinked leaves (NCL). Input and elution fractions were then analyzed by Western blots. FMT was detected in CL elution fractions with either FMT or GFP antibodies. ARGONAUTE 1 (AGO) and UGPAse were used respectively as positive and negative controls for oligo-dT pull down. L, ladder

### *fmt* knock-down (KO) is associated with reduced growth of seedlings and slightly affects the mitochondrial proteome

To identify FMT functions, an *fmt* KO mutant (SALK_056717) was analyzed. The T-DNA insertion in *the FMT* locus (AT3G52140) was verified (**Figure 6 A-B**), and FMT protein was not detected in this line (**Figure 6C**). *fmt* seedlings appeared smaller, with shorter primary roots than wild type (**Figure 6D**). The difference between *fmt* and Col0 was lost over time, and the delayed phenotype was no longer observed in older and flowering plants (**Figure 6E**). Last, the clustering of mitochondria was observed in the SALK_056717 line (**Figure 6F**), as it has been previously observed in ethyl methanesulfonate (EMS), SALK_046271, and SAIL_284_D06 *fmt* lines (Logan et al., 2003;El Zawily et al., 2014;Ayabe et al., 2021).

**Figure 6:**
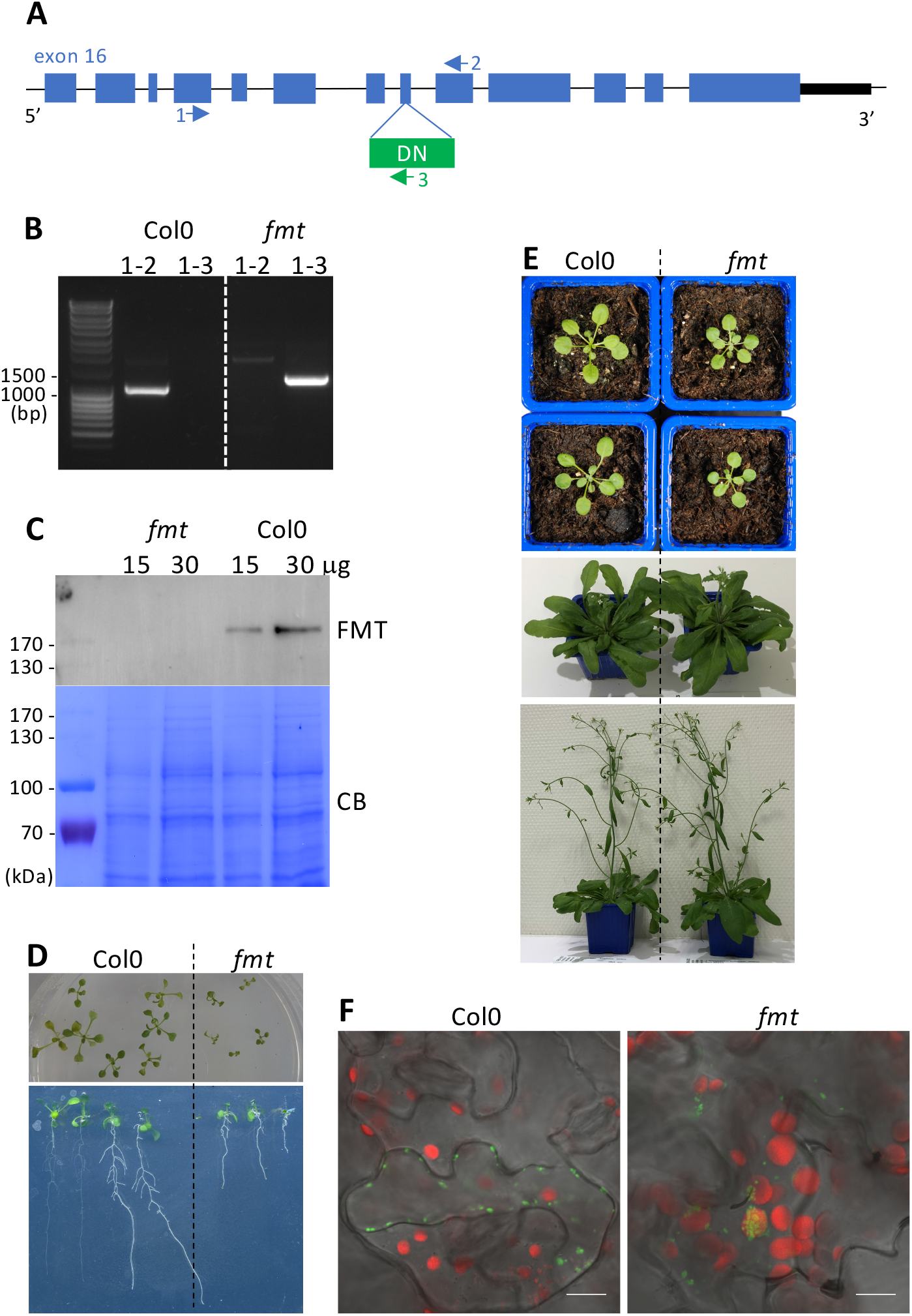
Characterization of *fmt* KO line SALK_056717. **A-** Schematic representation of T-DNA insertion in 3’ end of AT3G52140 locus. Blue boxes, exons; black box, 3’ untranslated region; arrows, primers used for PCR genotyping. **B**-PCR genotyping. Expected size of PCR products: 1060 bp in Col0 with primers 1-2, and 1300 bp in *fmt* with primers 1-3. **C**-FMT protein level in total extracts from Col0 and *fmt* leaves. Western blot with anti-FMT antibody (FMT); CB Coomassie Blue staining of the membrane. **D**-*fmt* phenotype on agar plate (13-day-old) **E-** *fmt* phenotype on soil (3, 8 and 10-week-old). **F**-mitochondria phenotype. *At* leaves transiently expressing COX IV-GFP (mitochondrial protein, in green) were examined by confocal microscopy. Red: chloroplast autofluorescence. Scale bar: 10 m

According to *fmt* seedlings phenotype (**Figure 6D**), seedlings mitochondria were deeper analyzed. The DNA, RNA and proteins contents were analyzed. The quantity of mitochondrial DNA in a total DNA extract appeared similar in Col0 and *fmt* (**Figure 7A**). Moreover, a global effect on mitochondrial-encoded RNAs was not observed in *the fmt* line (**Figure 7B**), even if some RNAs appeared weakly affected (COX1, RPS3, 18S). We also compared mitochondrial proteomes in seedlings of *fmt* and Col0. Mitochondrial extracts were analyzed by mass spectrometry. About 720 mitochondrial proteins were identified in each extract, and 72 proteins were enriched or depleted in *fmt* mitochondria compared to Col0 (**Figure 7C**; **Table S5**; **Figure S5A**). Eleven were over-expressed among which the 3 alternatives oxidases, which is consistent with AOX activity and Western blot (see below, **Figure 7D-E**). Among the 61 down-regulated proteins, 3 were TCA enzymes, 6 were components of the mitochondrial ribosome (Waltz et al., 2019) and 22 were from OXPHOS complexes, mostly from complex I (14 out of the 22) (Senkler et al., 2017). The lower level of some mito-ribosomal proteins is consistent with the lower level of the mitochondrial 18S rRNA (**Figure 7B**).

**Figure 7:**
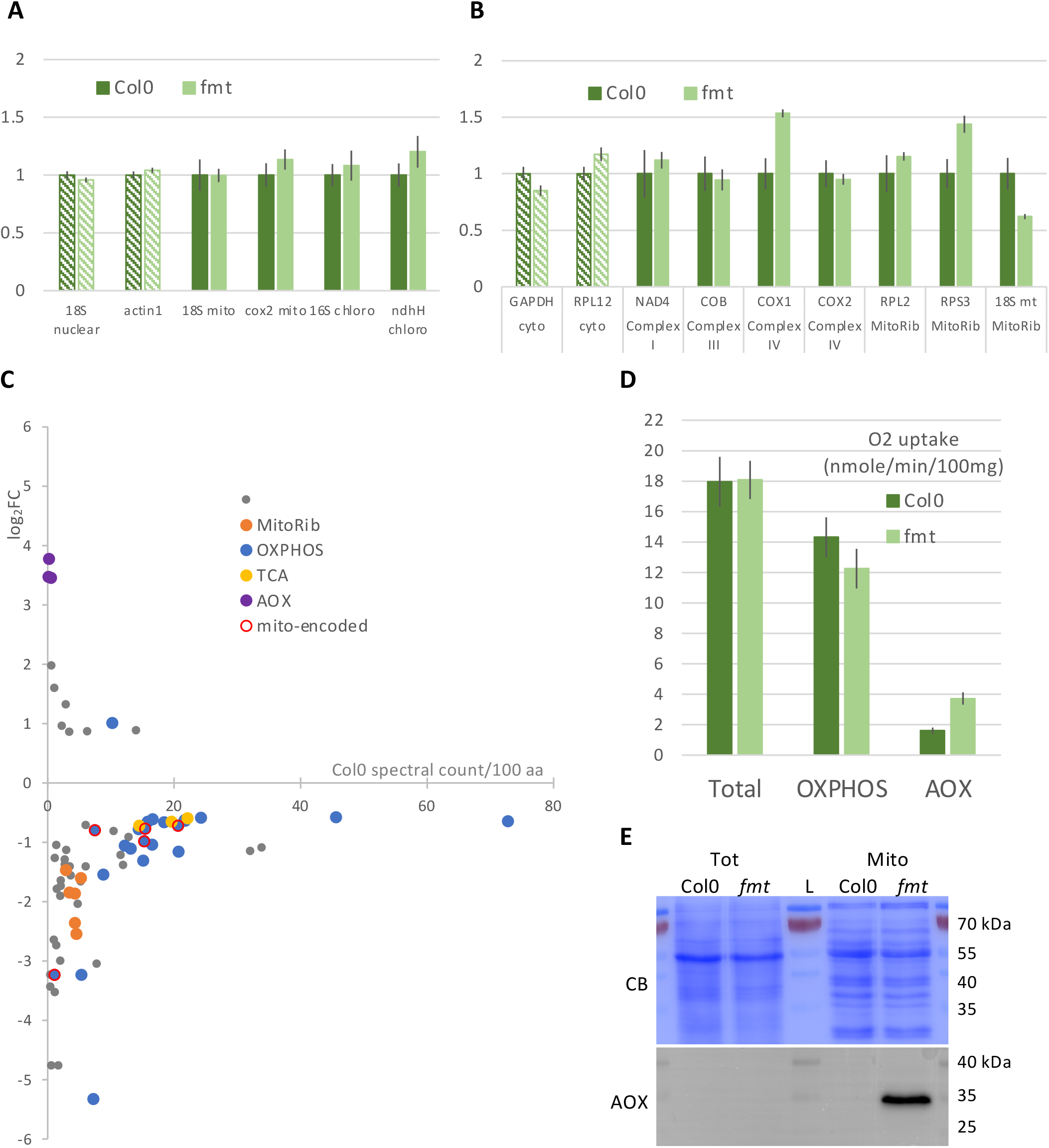
Analysis of mitochondria from *fmt* seedlings. **A-** Mitochondrial DNA content in total extracts. The level of organellar DNA was evaluated in total extracts of Col0 or *fmt* seedlings by qPCR. Normalisation was done with 2 nuclear genes coding for cytosolic 18S and actin, and results are expressed relatively to Col0. The error bars correspond to SEM (biological repeats n=3). **B-** Mitochondrial RNA level in total extracts. The level of mitochondrial encoded RNAs was evaluated in total extracts of Col0 or *fmt* seedlings by RT-qPCR. Normalisation was done with 2 cytosolic transcripts (GAPDH and RPL12), and results are expressed relatively to Col0. The error bars correspond to SEM (biological repeats n=3). **C-** Mitochondrial proteome. Proteins from purified mitochondria were analysed by MS/MS, and spectral count label-free quantifications were performed (*fmt* compared to Col0). With criteria [adjusted p-value < 0.1, Fold Change > 1.5], 72 were found differently expressed in *fmt* mitochondria compared to Col0. The expression level in Col0 is presented in the x-axe (spectral count per 100 amino acids). The fold change is shown in y-axe (*fmt* versus Col0)). **D**-O_2_ uptake (nmol/min/100 mg fresh seedlings): OXPHOS and AOX activities. Oxygen consumption was measured with an oxygraph. The addition of cyanide and of propygallate allowed to evaluate OXPHOS respiration and AOX activity (see Fig S6) (n=10). **E**-Western blot of total and mitochondrial extracts from 8-day-old *fmt* and Col0 seedlings. Immunodetection was performed with antibodies against the mitochondrial AOX. CB, Coomassie Blue staining of the membrane.

A similar approach was performed with mitochondria from *fmt* and Col0 inflorescences. With the same criteria as for seedlings, only 4 proteins were found affected in *fmt* mitochondria (**Figure S5B, Table S5**), all 4 being also affected in *fmt* seedlings mitochondria. Such a result is consistent with the absence phenotype in flowering plants.

Oxygen uptake was also measured in Col0 and *fmt* seedlings (**Figure 7D**), but no difference was observed between the 2 lines. The addition of cyanide (KCN), which blocks oxidative phosphorylation (OXPHOS pathway), and propyl gallate (PG), which blocks alternative oxidase (AOX), showed an increase of AOX pathway but no apparent change in OXPHOS.

In summary, the reduced levels of some OXPHOS proteins were observed in seedling mitochondria, but they did not seem to affect significantly OXPHOS activity. For mitochondrial translation, some mito-ribosomal proteins were also found down-expressed in *fmt*. The proteome analysis showed that most of the 16 identified mitochondrial-encoded proteins seemed weakly depleted in *fmt* (**Figure S5C**), suggesting that the mitochondrial translation could be weakly affected in this mutant. Last, increased levels of AOX proteins and AOX activity were observed in *fmt*. AOX have been usually considered as indicators of stress and mitochondrial dysfunctions (Robert et al., 2012; Juszczuk et al., 2012; Saha et al., 2016).

### The association of cytosolic mRNAs with the mitochondrial surface is disrupted in *fmt* line

A microarray-based transcriptomic analysis has been previously performed in an EMS mutant of FMT (El Zawily et al., 2014), and only minor changes in mRNAs abundance were observed. Among the affected transcripts, some mRNAs coding for mitochondrial proteins have been identified as mitochondrial-associated in *Solanum tuberosum* (MLR RNAs) (Vincent et al., 2017). We have explored the expression and mitochondrial association of 3 such mRNAs: CIII-RISP (AT5G13430), coding for the RISP subunit in complex III (Li et al., 2019), and 2 complex I subunits, CI-51 (AT5G08530) and CI-7.5 (AT1G67785) (Klodmann and Braun, 2011).

In addition, in *A. thaliana*, a well-studied targeted mRNA is VDAC3 (AT5G15090). VDAC3 gene has been previously shown to be transcribed into 2 isoforms that differ by the length of their 3’ UTR. The long isoform was found at the surface of mitochondria, but not the short one (Michaud et al., 2014).

To test the impact of *fmt* mutation on these mRNAs, total and mitochondrial RNAs were extracted from Col0 and *fmt* seedlings, and RT-qPCR were performed. CI-51 appeared down expressed in *fmt* total extracts, but neither CI-7.5, nor CIII-RISP. In contrast, VDAC3-long appeared overexpressed in *fmt*, which is consistent with the fact that this isoform is highly sensitive to stress (Hemono et al., 2020) (**Figure 8A**).

**Figure 8:**
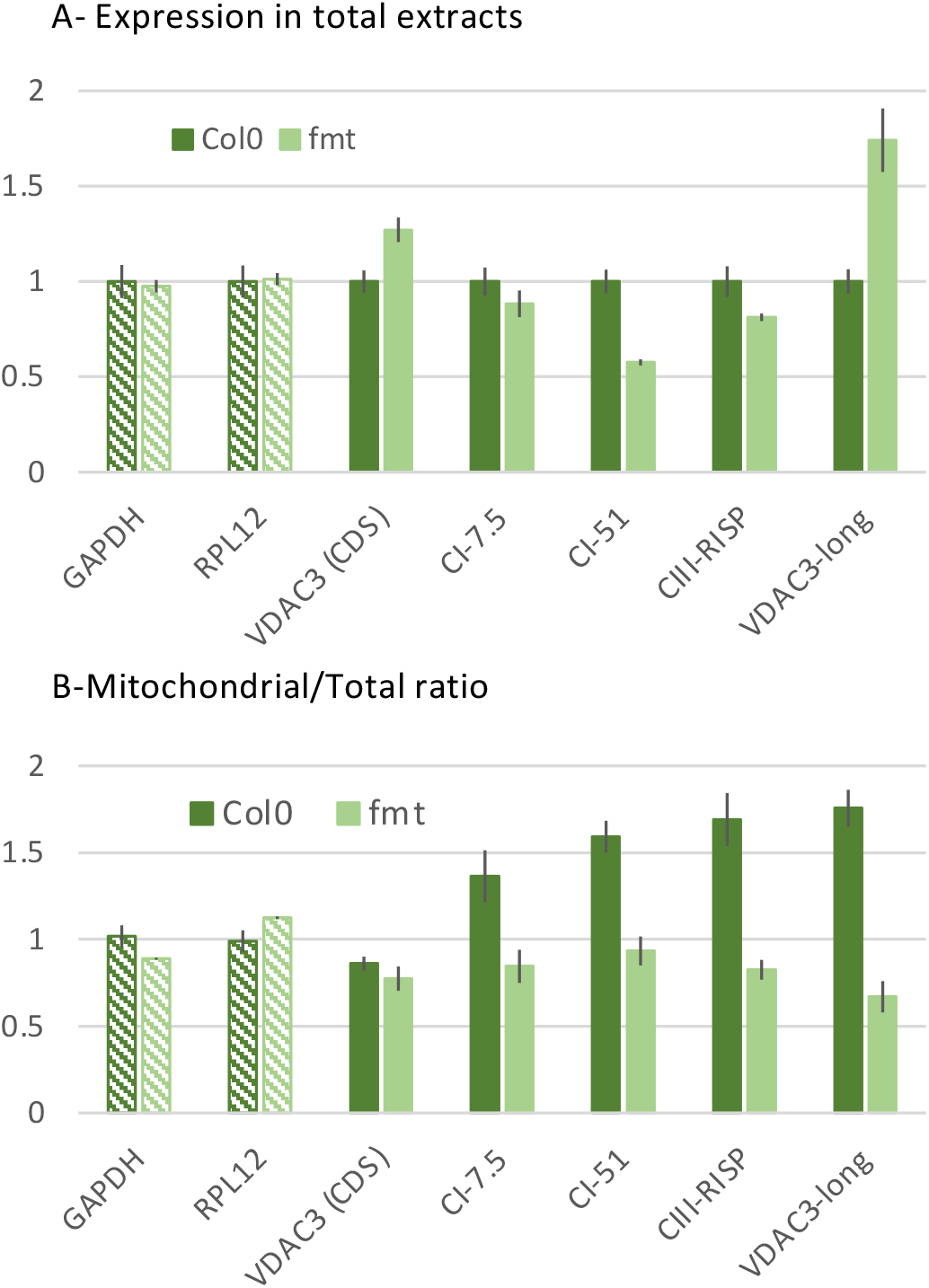
*fmt* mutation disturbs mRNA targeting to the mitochondrial surface. Messenger RNAs were quantified by RT-qPCR in mitochondrial and total extracts from seedlings. To compare the 3 biological replicates, normalization was performed with GAPDH and RPL12 mRNAs. **A**-Total extracts: The quantity of each mRNA in *fmt* total extracts was expressed relative to its level in wild-type Col0**. B**-Mitochondrial/Total ratio. The quantity of each mRNA in the mitochondrial fraction was corrected by its quantity in the total fraction to take into account the expression level of the gene. Mitochondrial/Total ratios were expressed relative to the mean of GAPDH and RPL12 ratios. Error bars represent SEM (n = 3). GAPDH, glyceraldehyde-3-phosphate dehydrogenase (AT1G13440); RPL12, cytosolic ribosomal protein L12A (AT2G37190); VDAC3, mitochondrial voltage-dependent anion channel 3 (AT5G15090) (VDAC3 (CDS) corresponds to both VDAC3 mRNAs isoforms); CIII-RISP, RISP subunit in complex III (AT5G13430); CI-51 (AT5G08530) and CI-7.5 (AT1G67785), 51kDa and 7.5 kDa proteins in complex I.

As already shown (Michaud et al., 2014), VDAC short and long isoforms differ in their association with mitochondria. The 3 other mRNAs were found enriched in the Col0 mitochondrial fraction compared to VDAC3 short isoform. However, in the *fmt* line, the mitochondrial associations of the 4 mRNAs appeared similar to the non-targeted VDAC3 short isoform mRNA (**Figure 8B**), suggesting a role of FMT protein in mRNA association with the mitochondrial surface.

CIII-RISP and CI-51 proteins were identified in the mitochondrial proteome, and both were among the 61 down-regulated proteins in *fmt* (**Figure 7C**). The steady-state of CIII-RISP protein dropped while the steady-state of its mRNA was not clearly affected, suggesting a correlation between mRNA localization and protein abundance in mitochondria. The decrease in CI-51 protein could be explained by the decrease in its mRNA level and/or in its mRNA association with mitochondria. For CI-7.5 and VDAC3 proteins, no hypothesis could be made, since CI-7.5 was not detected in the proteome, and 2 mRNAs with different behaviors code for the same VDAC3 protein.

## DISCUSSION

FMT is an evolutionarily conserved protein belonging to the CLU family, and found in species as distant as plants, yeast, amoeba, flies, or mammals. We have shown that FMT is a cytosolic RBP, sometimes found near the mitochondria. By coIP and reverse coIP experiments, we have found that FMT copurifies with the cytosolic ribosome, but only in mitochondrial fractions. We have confirmed the proximity of FMT with the cytosolic ribosome and with the mitochondria by split-GFP experiments.

As with all proteins of the CLU family, the *fmt* mutation-induced clustering of the mitochondria. In plants, *fmt* mutation was also found to have a decreased capacity to fight bacterial infection (Vellosillo et al., 2013). The consequences of the mutation on plants grown in standard conditions were evident at the level of seedlings, with growth retardation, but this phenotype faded during development. At the mitochondria level, the mitochondrial proteome from seedlings appeared to be affected, with a decrease in the abundance of OXPHOS proteins and components of the mitoribosome. On the other hand, AOX proteins were overexpressed, suggesting a mitochondrial stress. The dysfunction of the mitochondria in young plants is confirmed by the high proportion of depolarized mitochondria in *fmt* mutant (Nakamura et al., 2020; Ma et al., 2021). However, the defect in the mitochondrial proteome was no longer visible in inflorescences, in agreement with the absence of phenotype in flowering plants. So, the function of FMT appears particularly important in the early stages of development.

Mitochondrial biogenesis relies on the massive import of proteins encoded by the nuclear genome. This import can occur post-translationally or co-translationally. In this last case, the mRNAs are sent to the surface of the mitochondria, where cytosolic ribosomes translate them, and the proteins are directly imported into the mitochondria. Finding FMT associated with cytosolic ribosomes in mitochondrial fractions prompted us to explore the co-translational import pathway. This pathway is still little studied in *A. thaliana*, and a single mRNA, VDAC3-long, has so far been identified on the surface of mitochondria (Michaud et al., 2014). Therefore, we looked for other potential candidates among the mRNAs whose expression is reduced in *fmt* mutants (El Zawily et al., 2014). We have thus selected 3 mRNAs whose orthologs in *Solanum tuberosum* are targeted to the mitochondrial surface (Vincent et al., 2017). These 3 mRNAs and VDAC3-long were no longer enriched at the mitochondrial surface in *fmt* mutant, showing the implication of FMT in the targeting and/or anchoring of these mRNAs to the mitochondrial surface.

The localization of an mRNA on the mitochondrial surface requires its transport from the nucleus to the mitochondria and its docking to the mitochondria, where it would be translated. These mechanisms are still poorly understood, with the intervention of multiple signals (on the mRNA and/or on the protein) and various protein factors. FMT is an abundant RBP in the cytosol. It is also found associated with ribosomes on the mitochondrial surface. FMT could therefore be involved, directly or indirectly, in the targeting of its target mRNAs, their anchoring on mitochondria, or their localized translation. This role should be further explored as part of the latest discoveries on FMT, which has recently been shown to be associated with mitophagy processes. Ma *et al.* have shown that treatment with a decoupling agent induces mitophagy and a reorganization of FMT in foci, some of which co-localize with the mitochondria. In addition, FMT has been shown to associate with ATG8, a mitophagosome marker (Ma et al., 2021). Nakamura *et al.* have shown that the clustering of the mitochondria was higher in an *atg5/fmt* double mutant than in a *fmt* mutant (ATG5 is also involved in mitophagy) (Nakamura et al., 2020). Therefore, the relationship between FMT and mitophagy needs to be further explored. Finally, a last point to be addressed concerns the link between FMT and chloroplasts. Photosystem I activity is reduced in *fmt* (El Zawily et al., 2014). Moreover, the de-etiolation of seedlings germinated under dark conditions is reduced in *fmt* and also in *atg5*. These results suggest that the role of FMT could be not limited to the mitochondria but would also impact inter-organellar communications and consequently the functioning of the plant cell in response to development or biotic or abiotic signals.

## METHODS

### Plant material and growth conditions

*A. thaliana* plants were grown in long-day conditions (16 h-day-at 21°C / 8 h night at 18°C cycles, LED tubes Philips 1500 mm SO 20W 840 T8, photon flux density of 120 μmol/s/m^2^ at the plant level). Seedlings were grown for 8 days in hydroponic cultures at 23°C with constant light in Murashige and Skoog MS231 medium (Duchefa).

The *FMT* insertion mutant SALK_056717 was from Columbia ecotype. The line expressing the epitope-tagged ribosomal protein, FLAG-RPL18 (AT3G05590), was from (Zanetti et al., 2005). FMT-GFP, GFP-FMT and MS2-GFP lines were obtained by floral dip with pAM557, pAM549 and pAM495 plasmids respectively (**Table S6**) {Clough, 1998 #958}.

Transient transformation of *At* and *N. benthamiana* was performed by infiltration of leaves with a suspension of *Agrobacterium tumefaciens* harboring different constructs. The constructs allowed the expression of different GFP or split-GFP constructs, the pSu9**–**RFP mitochondrial marker (Michaud et al., 2014), the COXIV-GFP mitochondrial marker (Peeters et al., 2000) and the silencing suppressor P19 protein (Michaud et al., 2014).

### Genotyping

Genomic DNA was extracted from rosette leaves of 4-week-old *At* plants as described in {Edwards, 1991 #959}) and subjected to a PCR-based screening using primer pairs 1-2 and 1-3 (1 : 5′ CAACCCATCACCAAGAAGGGTC 3′; 2 : 5′ CATGGTACAAAACCATAGCCAGG 3′; 3 : 5′ TGGTTCACGTAGTGGGCCATCG 3′). PCR products were separated by agarose gel electrophoresis and visualized by ethidium bromide staining.

### qPCR

According to the manufacter’s instructions, genomic DNA for qPCR was extracted from 8-day-old water-cultured seedlings using NucleoSpin® plant L kit. For qPCR, three biological replicates were performed. DNA was used at 3 ng/μL and 10 times diluted, and 3 technical replicates were performed for each dilution. The qPCR efficiency for each primers couple was determined by dilutions of DNA. The qPCR results were normalized with the nuclear genes coding for actin (Actin cyto) and 18S ribosomal RNA (18S cyto) (Table S7) to compare the different replicates.

### RNA extraction, reverse transcription and RT-qPCR

RNA was extracted from mitochondria and whole cells using Tri Reagent® (Molecular Research Center) according to the manufacter’s instructions, then treated with RNase free-DNase RQ1 (Promega) and quantified with nanodrop. Their quality was checked by electrophoresis in MOPS buffer/formaldehyde/agarose gel. Reverse transcription was performed with Reverse Transcription SuperScript™ IV (Invitrogen) in presence of hexamers and oligo dT primer (Michaud et al., 2010). For qPCR, the RT was used directly and 10 times diluted, and 2 to 3 technical replicates were performed for each dilution. Three biological replicates, corresponding to mitochondrial and total RNA extracted from the same plant material, were prepared. The qPCR efficiency for each primers pair was determined by dilutions of cDNA. To compare the different replicates, the qPCR results were normalized with the cytosolic ribosomal protein L12 (RPL12, AT2G37190) and glyceraldehyde-3-P-dehydrogenase (GAPDH, AT1G13440) mRNAs.

### Mitochondria preparation

Gradient-purified mitochondria were prepared from 8-day-old water-cultured seedlings according to {Hemono, 2020 #954}, or from inflorescences of 6/8-week-old *Arabidopsis thaliana* according to {Waltz, 2019 #849}.

### Immunoprecipitation

Proteins were extracted from grounded seedlings or inflorescences or from corresponding mitochondria. μMACS™ GFP or μMACS™ DYKDDDDK (FLAG) Protein Isolation Kits (Miltenyi Biotec) were used for immunoprecipitations. Three biological repeats were performed. For seedling material, the composition of the lysis buffer was [50 mM Tris-HCl pH8, 50 mM NaCl, 10 mM MgCl2, 1% Triton X-100, 200μg/mL cycloheximide, protease inhibitors (cOmplete™, EDTA-free protease inhibitor cocktail, Roche, 1 tablet /50 mL)]. The wash buffer corresponded to the lysis buffer but with 0.1% Triton X-100. For inflorescence material, the composition of the lysis buffer was [20 mM HEPES-KOH pH7.6, 100 mM KCl, 20 mM MgCl2, 1 mM DTT, 1% Triton X-100, 20μg/mL cycloheximide, protease inhibitors). The wash buffer corresponded to the lysis buffer but with 0.1% Triton X-100.

### Proteomic analyses

Proteins (1 mg) from each sample were trypsin digested, then analyzed by nanoLCMS/MS. Data were searched against the TAIR *A. thaliana* database. All these steps were performed at the “Plateforme Protéomique Strasbourg-Esplanade” (http://www-ibmc.u-strasbg.fr/proteo/Web/accueil.htm).

To identify significantly affected proteins, a statistical analysis based on spectral counts was performed using a homemade R package (https://github.com/hzuber67/IPinquiry4) as described in (Scheer et al., 2021). To avoid too much bias, only proteins with a mean of at least 3 spectra in the most expressed condition were further considered.

Mitochondrial localization was determined according to SUBA4 (http://suba.live/) (Hooper et al., 2017).

### Western blot and antibodies

Western blot analysis was conducted according to standard. Antibodies against AOX (T. Elthon, GT monoclonal antibodies, University of Nebraska, Lincoln, USA) (Elthon et al., 1989)), UGPase, Calnexin, AGO, and Arf1 (Agrisera), GFP and VDAC (A. Dietrich, IBMP, Strasbourg), were used. For the FMT antibody, the peptide corresponding to the C-terminal part of FMT (373 aminoacids) was purified from SDS-polyacrylamide gel and injected into rabbits (Covalab antibody production).

### Confocal microscopy

*N. bentamiana* leaves were observed 2 days after agroinfiltration*. A. thaliana* or *N. bentamiana* leaves were directly observed by using a LSM780 confocal microscope (Zeiss). GFP and RFP fluorophores were excited at 488 and 555 nm, respectively, and emission signals were simultaneously collected at 488–560 nm for GFP and at long-pass 560 nm for RFP.

### Oligo (dT) affinity purification of crosslinked protein–RNA complexes

Oligo(dT) capture was performed according to {Bach-Pages, 2020 #964} with minor modufications. For cross-linking, leaves of *At* GFP-FMT line were placed on ice and irradiated in a Stratalinker (Stratagene) with 254-nm UV light at 1000 mJ/cm2 (Cross-linked leaves, CL). The irradiation was performed twice with a 1-min pause in between treatments. After irradiation, leaves were immediately frozen in liquid N_2_ and ground into fine powder. Non-crosslinked leaves (NCL) were processed as a control. Leaves powder (1.1 g) was resuspended in 11ml lysis buffer (20 mM Tris HCl, pH 7.5, 500 mM LiCl, 0.5% LiDS, 1 mM EDTA, 0.2% IGEPAL, 2.5% PVP40 (wt/v), 1% B-ME (v/v), 5mM DTT, protease inhibitor and Vanadyl RNase inhibitor) and incubated for 10 min on a rotator at 4°C. The lysate was cleared by two centrifugations (4000 rpm, 10 min, 4 °C and 20000g, 10 min, 4°C) and filtered (Miracloth, Merck 475855-1R). The lysate (input) was then incubated with 200 μl oligo(dT)25 magnetic beads (New England Biolabs) for 1 h at 4°C on a rotator. Beads were collected on a magnet, washed twice with 1 mL lysis buffer, twice with 1mL buffer I (20 mM Tris HCl, pH 7.5, 500 mM LiCl, 0.1% LiDS, 1 mM EDTA and 5 mM DTT), twice with 1mL buffer II (20 mM Tris HCl, pH 7.5, 500 mM LiCl, 1 mM EDTA and 5 mM DTT), and twice with 1mL buffer III (20 mM Tris HCl, pH 7.5, 200mMLiCl, 1 mM EDTA, and 5 mM DTT). Beads were resuspended in 100 μl protein extraction buffer (8M Urea, 50mM Tris HCL pH 6.9, 1mM EDTA, 5% β-mercaptoethanol) and directly used for immunoblotting.

### O2 consumption

Oxygen consumption of seedlings was measured in 2 mL of culture medium with a liquid-phase Oxytherm oxygen electrode system (Hansatech Instruments, Pentney, UK). Five-day-old seedlings grown in liquid phase (30–40 mg) were directly imbibed in the electrode chamber and oxygen consumption rates were measured. After 6 min, KCN was added (final concentration: 2.5 mM), and oxygen consumption was re-measured. Differences between these two rates corresponded to cyanide-sensitive oxygen uptake (OXPHOS oxygen uptake). Then, after 8 min, propylgalate (PG) was added (final concentration: 2.5 mM) and oxygen consumption was re-measured. Differences between the KCN and the PG rates correspond to AOX oxygen uptake.

## Supporting information

Supplemental informations

## Authors contribution

Experiments: MH, TSG, JR, AV, PH, EU; Writing & Editing: PN, AMD

## SUPPORTING INFORMATION

**Figure S1: Quality control of proteins extracts.** Western blots were performed on total (Tot) and mitochondrial (Mito) extracts with antibodies against VDAC (30 kDa, mitochondria), UGPase (52 kDa; cytosol), Calnexin (67 kDa; ER), and Arf1 (21 kDa; Golgi). CB, Coomassie blue staining of the membrane. L, ladder.

**Figure S2: Identification of proteins co-purifying with FMT at the surface of seedlings mitochondria.** Co-immunoprecipitated proteins from GFP-FMT and Col0 mitochondrial samples were identified by LC-MS/MS. Only one replicate was performed. NSAF (Normalized spectral Abundance Factor) and Z-scores were determined for GFP-FMT and Col0 coIP (Paoletti et al., 2006; McIlwain et al., 2012). Log_2_FC (y-axis) was calculated with NSAF values, x-axis corresponded to Z-score in GFP-FMT experiment. Only proteins with a log_2_FC>2 are shown. Ribosomal proteins are in black, and mitochondrial ones are in orange.

**Figure S3: Identification of proteins co-purifying with FMT in total extracts from inflorescences (A), seedlings (B), and seedlings after formaldehyde crosslinking (C).** Only proteins with a mean of at least 3 spectra in FMT IP are shown (grey). The yellow spots correspond to the FMT bait. The enriched proteins are with a red circle (cuts-off FC> 2 and adj-p<0.1). In (C), proteins from cytosolic ribosomes (Salih et al., 2020) are indicated in black.

**Figure S4: Controls for Split-GFP experiments**. *N. benthamiana* leaves transiently expressing one protein fused with one part of GFP (blue, with GFP-β1-10; orange, with GFP-β11). White: Bright light; Green: GFP fluorescence; Red: chloroplast autofluorescence and mitochondrial pSU9-RFP fluorescence. All the scale bars correspond to 10 μm.

**Figure S5: Mitochondrial proteomes in seedlings and inflorescences** Proteins from purified mitochondria were analyzed by LC-MS/MS, and spectral count label-free quantifications were performed (*fmt* line compared to Col0). **A**-mitochondrial proteins from seedlings. **B**-mitochondrial proteins from inflorescences. **C**-mitochondrially encoded proteins in seedling mitochondria.

**Figure S6: O_2_ uptake (nmol/min/100 mg fresh seedlings):** Oxygen consumption was measured with an oxygraph. After a few minutes, cyanide (KCN), which blocks oxidative phosphorylation (OXPHOS pathway), was added. Then, a few minutes later, propylgallate (PG), which blocks alternative oxidase (AOX), was also added. The difference between total rate and residual rate in presence of KCN corresponded to OXPHOS respiration. The difference between rates in the presence of KCN and in the presence of both KCN and PG corresponded to AOX activity (Figure 7D).

**Table S1: Enriched proteins in co-immunoprecipitations from FLAG-RPL18 mitochondria.** Statistical analyses based on specific spectral counts identified 63 proteins enriched in IP from FLAG-RPL18 compared to both IP from FLAG-RPL18 total and Col0 mitochondrial extracts (cuts-off FC> 2 and adj-p<0.1).

**Table S2: Enriched proteins in FMT co-immunoprecipitations from inflorescences mitochondrial extracts.** Statistical analyses based on specific spectral counts identified 75 proteins enriched in IP from mitochondria of at least one FMT/GFP line compared to Col0 IPs (cuts-off FC> 2 and adj-p<0.1).

**Table S3: Enriched proteins in FMT co-immunoprecipitation from seedling mitochondrial extracts.** One hundred and twenty-four proteins were enriched in GFP-FMT compared to MS2-GFP IPs, with a FC above 2, among them 51 were cytosolic ribosomal proteins

**Table S4: Enriched proteins in FMT co-immunoprecipitation from seedling total extracts after formaldehyde crosslinking.** Statistical analyses based on specific spectral counts identified 119 proteins enriched in IP from crosslinked seedlings total extracts of FMT-GFP line compared to MS2-GFP IPs (cuts-off FC> 2 and adj-p<0.1).

**Table S5: Comparison of mitochondrial proteome in *fmt* and Col0 seedlings.** With a Fold-Change cut-off above 1.5 and an adjusted p-value below 0.1, 72 proteins were found affected in *fmt* compared to Col0

**Table S6: DNA constructs and plasmids.**

**Table S7: oligonucleotides used in qPCR and RT-qPCR**

