## Supplemental informations for "FRIENDLY (FMT) is an RNA binding protein associated with cytosolic ribosomes at the mitochondrial surface"

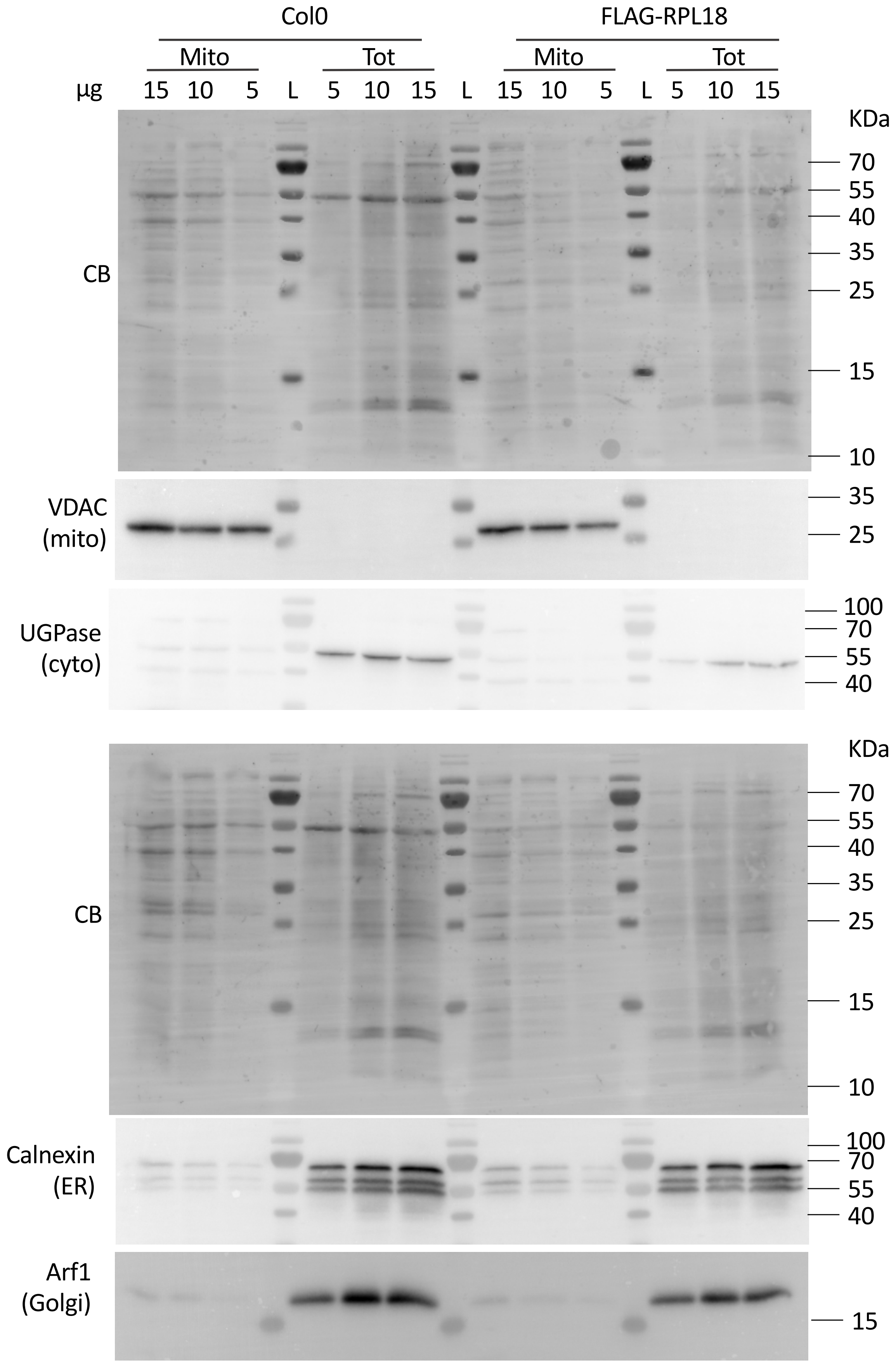


**Figure S1: Quality control of proteins extracts.**

Western blots were performed on total (Tot) and mitochondrial (Mito) extracts with antibodies against VDAC (30 kDa, mitochondria), UGPase (52 kDa; cytosol), Calnexin (67 kDa; ER), and Arf1 (21 kDa; Golgi). CB, Coomassie blue staining of the membrane. L, ladder.

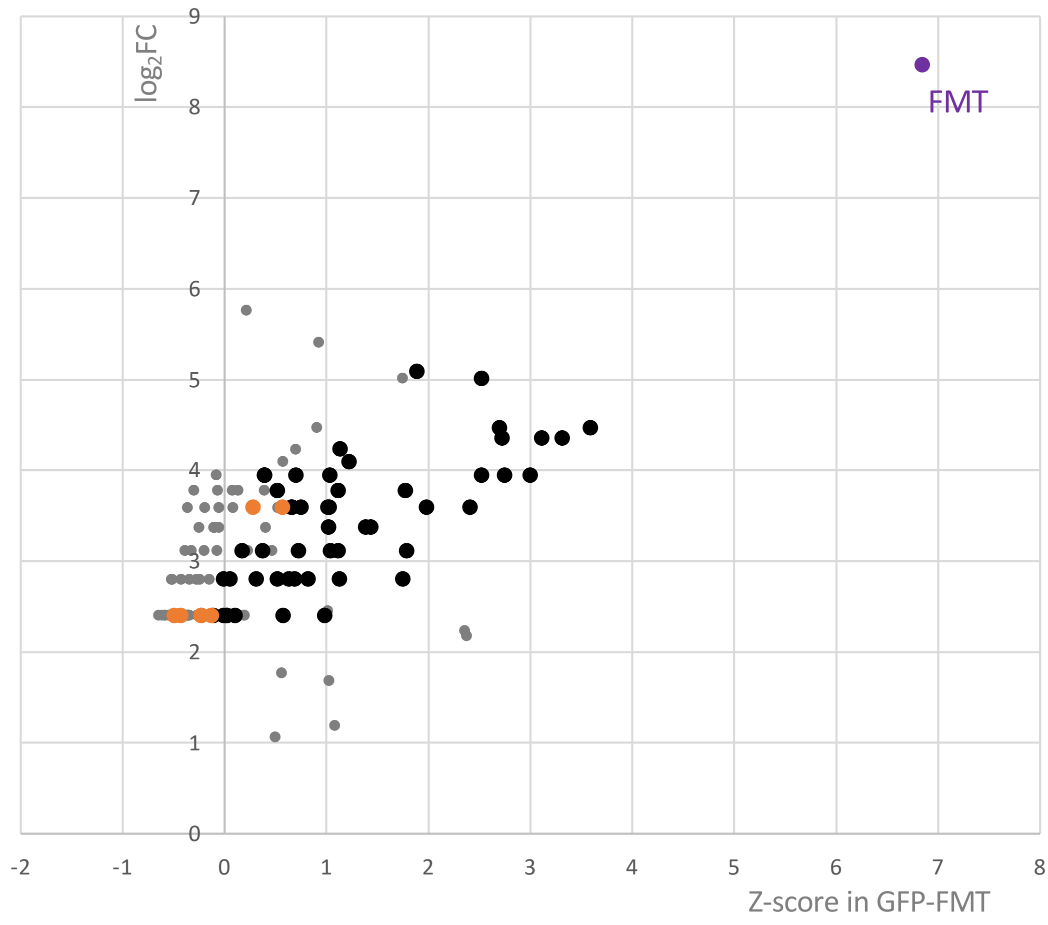


**Figure S2: Identification of proteins co-purifying with FMT at the surface of seedlings mitochondria.**

Co-immunoprecipitated proteins from GFP-FMT and Col0 mitochondrial samples were identified by LC-MS/MS. Only one replicate was performed. NSAF (Normalized spectral Abundance Factor) and Z-scores were determined for GFP-FMT and Col0 coIP (Paoletti et al., 2006) (McIlwain et al., 2012). $Log_2FC$ (y-axis) was calculated with NSAF values, x-axis corresponded to Z-score in GFP-FMT experiment. Only proteins with a $log_2FC>2$ are shown. Ribosomal proteins are in black, and mitochondrial ones are in orange.

**Figure S3: Identification of proteins co-purifying with FMT in total extracts from inflorescences (A), seedlings (B), and seedlings after formaldehyde crosslinking (C).** Only proteins with a mean of at least 3 spectra in FMT IP are shown (grey). The yellow spots correspond to the FMT bait. The enriched proteins are with a red circle (cuts-off FC> 2 and adj-p<0.1). In (C), proteins from cytosolic ribosomes (Salih et al., 2020) are indicated in black.

**A- from inflorescences** (control: Col0)

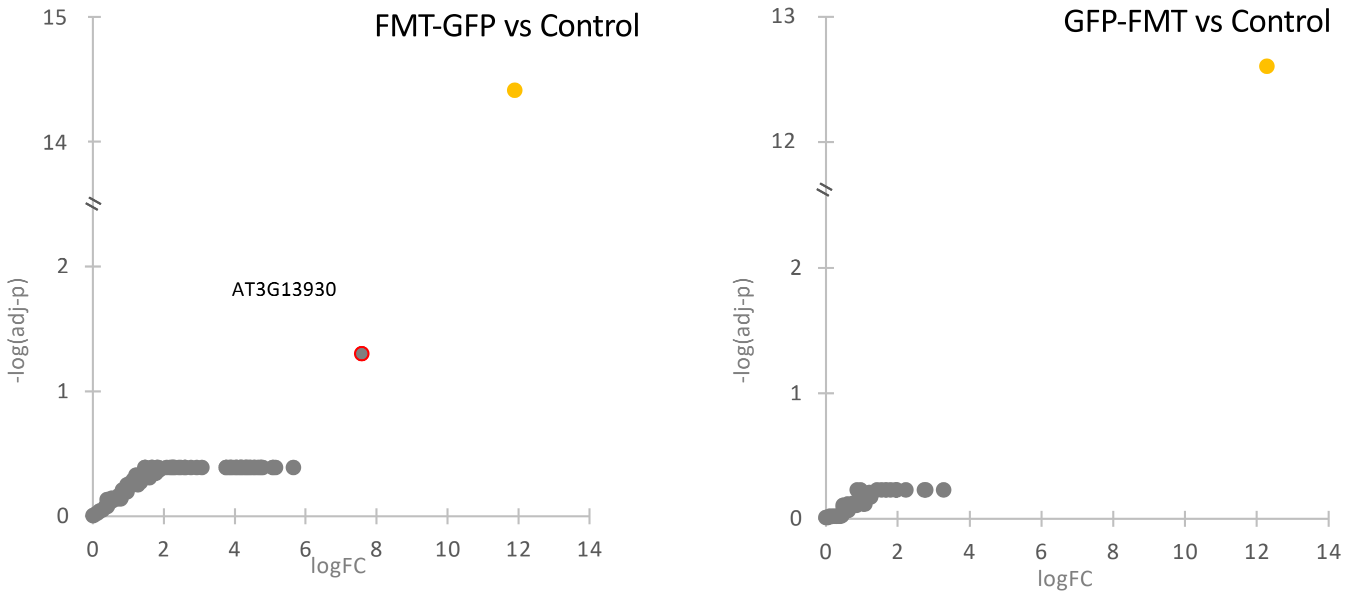


**B- from seedlings** (control: MS2-GFP)

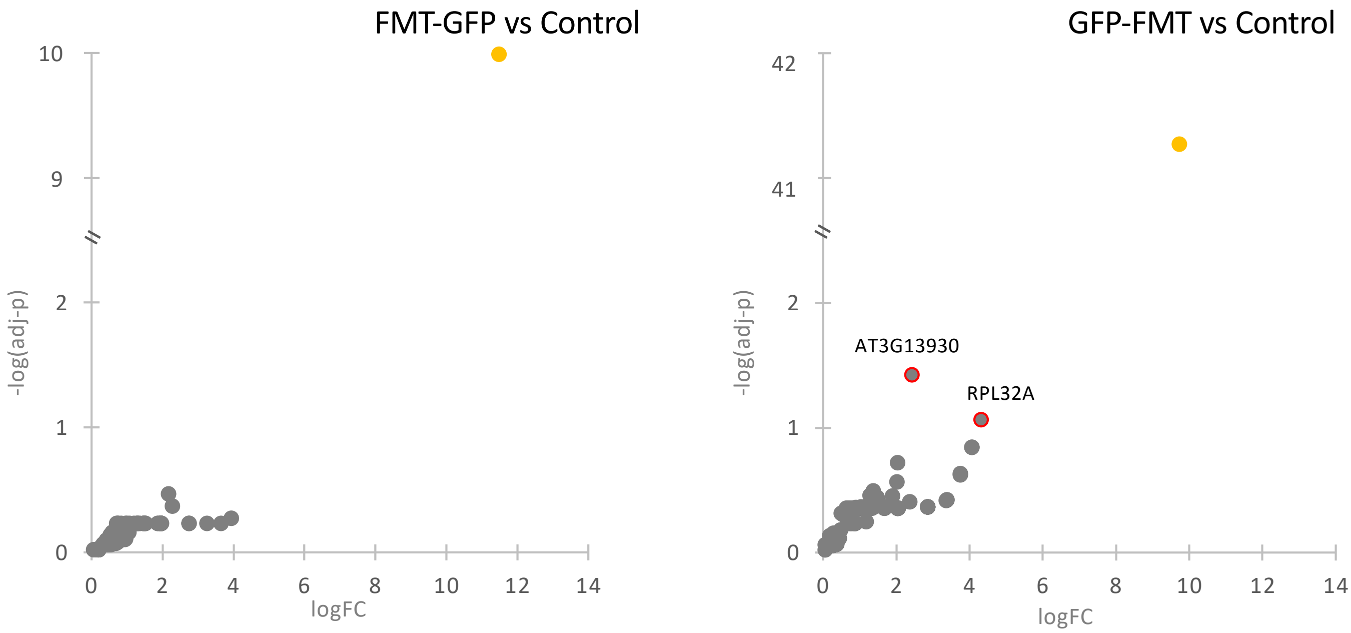


**C- from formaldehyde-crosslinked seedlings** (control: MS2-GFP)

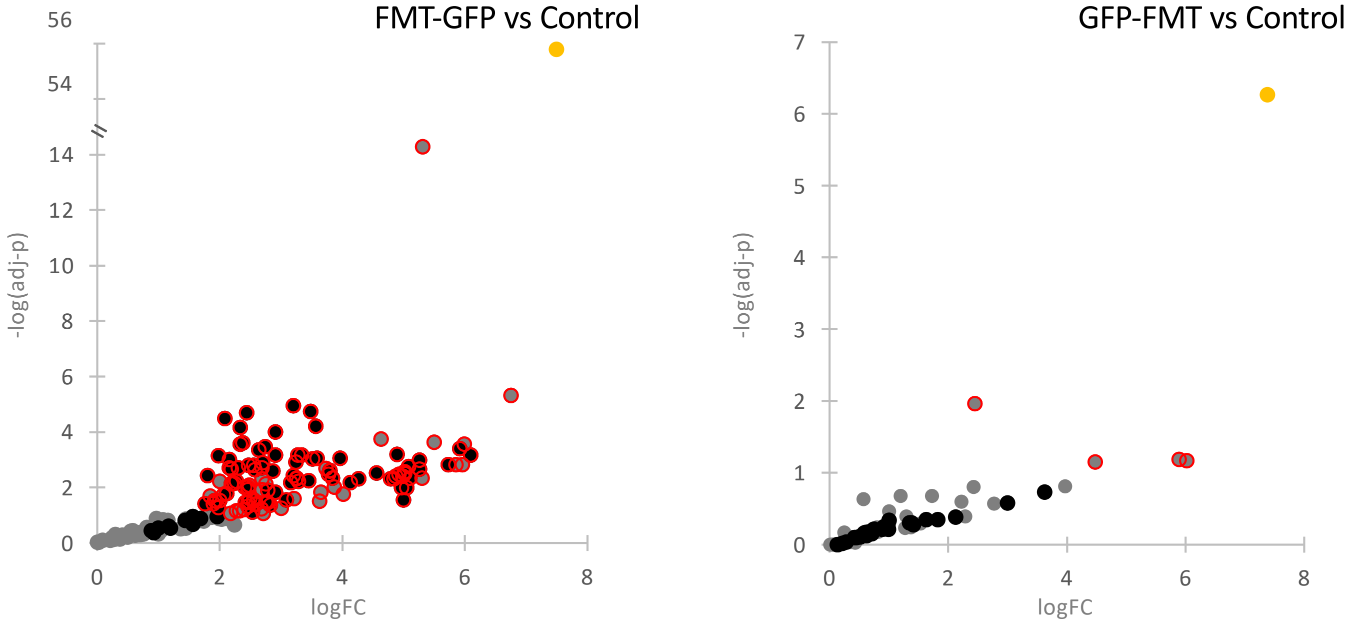

**Figure S4: Controls for Split-GFP experiments**. *N. benthamiana* leaves transiently expressing one protein fused with one part of GFP (blue, with GFP-β1-10; orange, with GFP-β11). White: Bright light; Green: GFP fluorescence; Red: chloroplast autofluorescence and mitochondrial pSU9-RFP fluorescence. All the scale bars correspond to 10 mm.

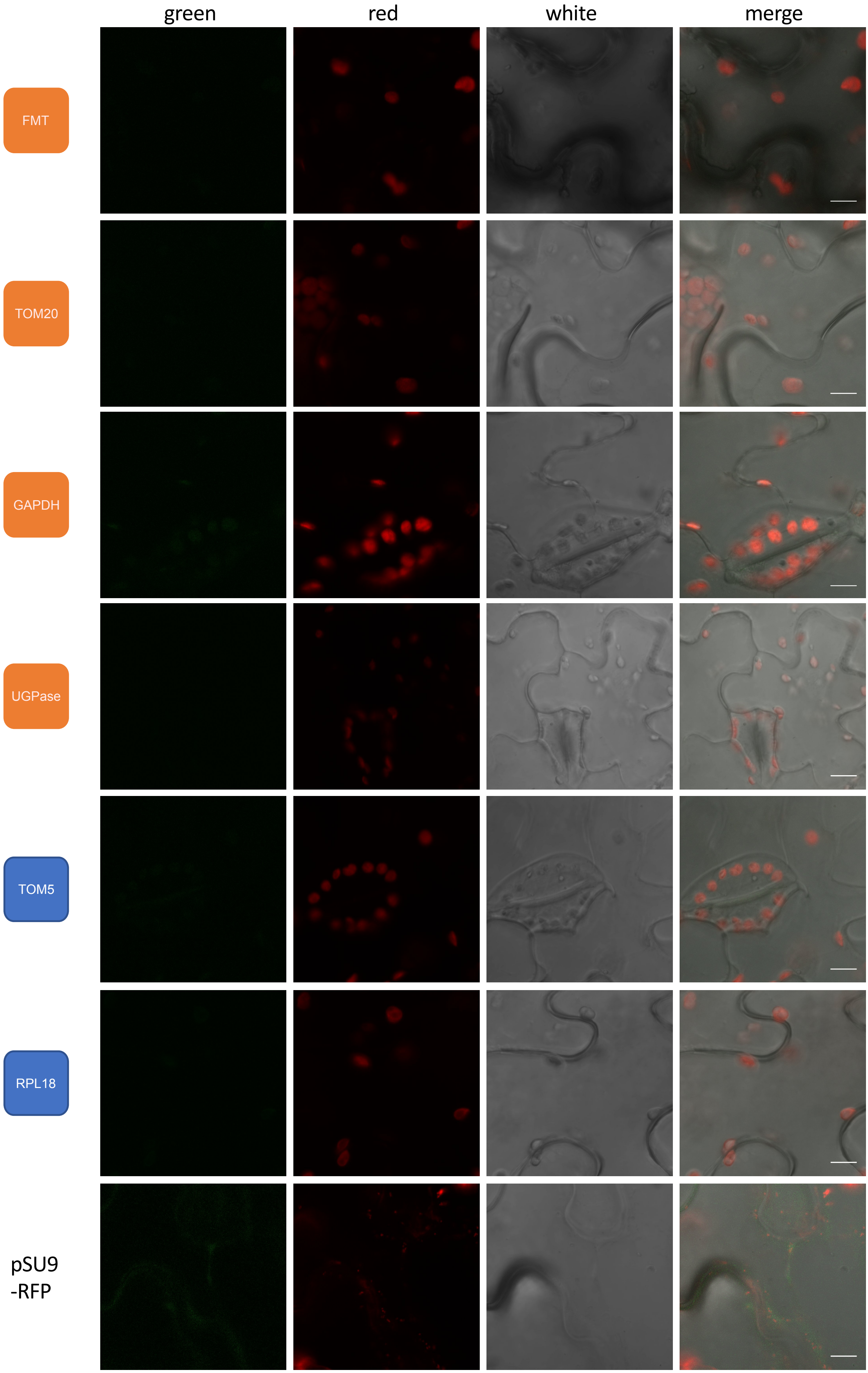

**Figure S5: Mitochondrial proteomes in seedlings and inflorescences.**
Proteins from purified mitochondria were analyzed by LC-MS/MS, and spectral count label-free quantifications were performed (*fmt* line compared to Col0). **A**-mitochondrial proteins from seedlings. **B**- mitochondrial proteins from inflorescences. **C**- mitochondrially encoded proteins in seedling mitochondria.

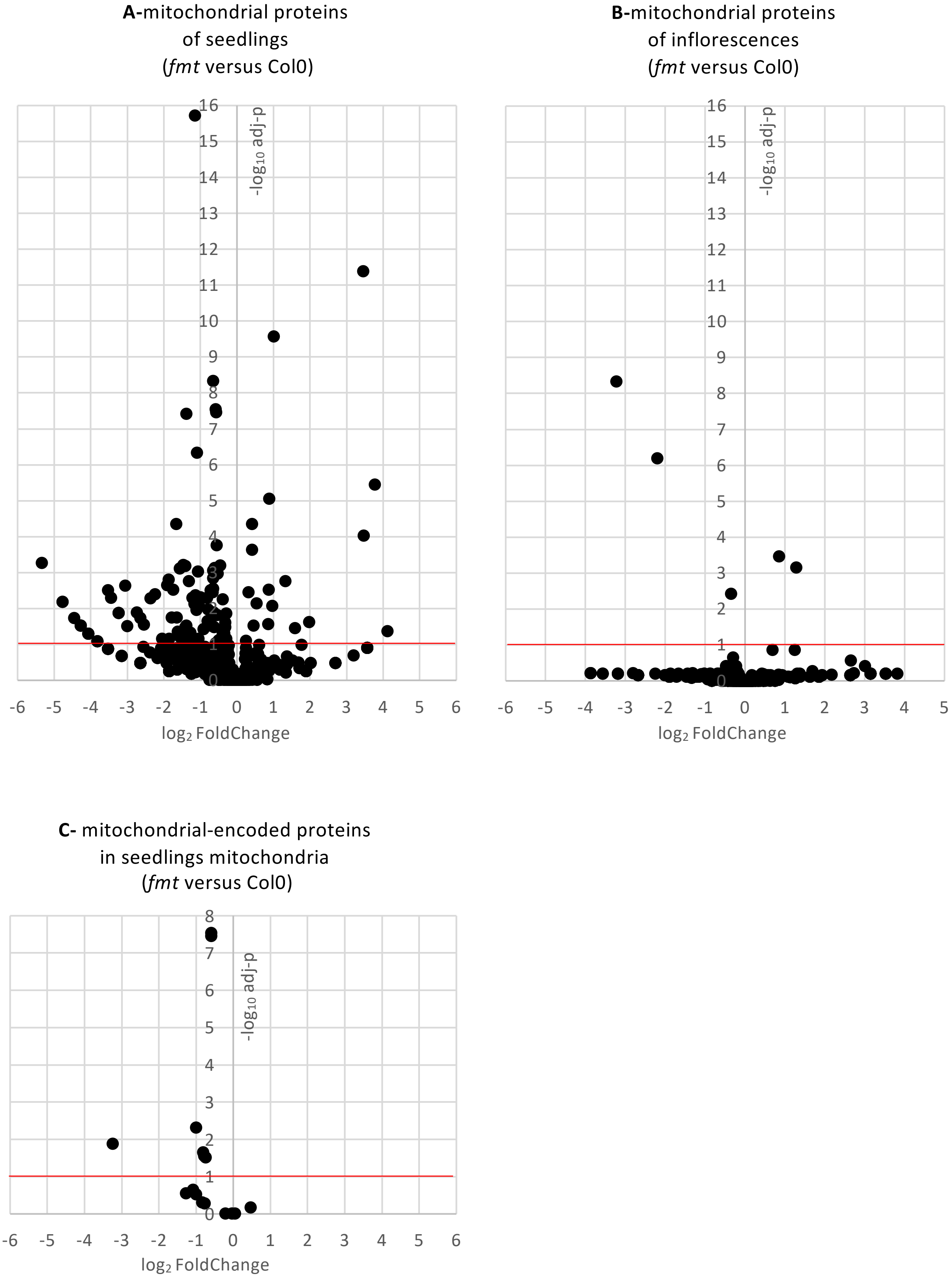

**Figure S6: $O_2$ uptake (nmol/min/100 mg fresh seedlings).**

Oxygen consumption was measured with an oxygraph. After a few minutes, cyanide (KCN), which blocks oxidative phosphorylation (OXPHOS pathway), was added. Then, a few minutes later, propylgallate (PG), which blocks alternative oxidase (AOX), was also added. The difference between total rate and residual rate in presence of KCN corresponded to OXPHOS respiration. The difference between rates in the presence of KCN and in the presence of both KCN and PG corresponded to AOX activity (Figure 7D).

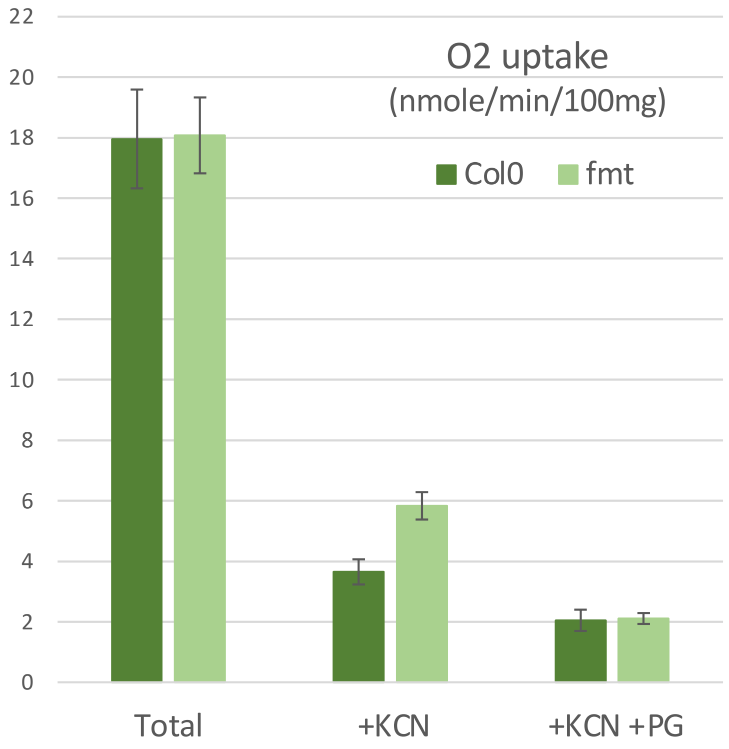

**Table S1**: Enriched proteins in co-immunoprecipitations from FLAG-RPL18 mitochondria. Statistical analyses based on specific spectral counts identified 63 proteins enriched in IP from FLAG-RPL18 compared to both IP from FLAG-RPL18 total and Col0 mitochondrial extracts

| AGI | Flag-RPL18 Mito_vs_Tot logFC | Flag-RPL18 Mito_vs_Tot -logadjp | Mito Flag-RPL18_vs_Col0 logFC | Mito Flag-RPL18_vs_Col0 -logadjp | description | name | cyto Rib (Salih 2020) | mito Rib (Waltz 2019) | SUBA4 |
|---|---|---|---|---|---|---|---|---|---|
| AT3G13300 | 7.40073545 | 9.01055018 | 5.57792627 | 9.06770808 | Transducin/V | VCS | | | cytosol |
| AT3G52140 | 6.95671841 | 7.39040559 | 6.98062727 | 8.44835425 | tetratricopep | FMT | | | cytosol |
| AT4G27740 | 6.92537171 | 7.30891851 | 1.91787736 | 2.6643708 | Yippee famil | | | | mitochondrion |
| AT5G19760 | 6.91979406 | 7.14874165 | 2.24685952 | 2.64716046 | Mitochondria | DTC | | | mitochondrion |
| AT4G29130 | 6.36489797 | 5.18243463 | 3.81278101 | 3.86251879 | hexokinase 1 | HXK1 | | | mitochondrion |
| AT1G78570 | 6.23933314 | 4.33254705 | 3.66771523 | 3.11944227 | rhamnose bi | RHM1 | | | cytosol |
| AT3G22310 | 6.21160158 | 4.9788107 | 1.63463488 | 1.11458662 | putative mit | RH9 | | | mitochondrion |
| AT4G01100 | 6.15786418 | 4.75696195 | 2.27681327 | 1.49842592 | adenine nucl | ADNT1 | | | mitochondrion |
| AT3G25150 | 5.99654342 | 4.42365865 | 6.02236768 | 5.00678839 | Nuclear trans | | | | nucleus |
| AT1G26630 | 5.99463665 | 4.39685563 | 2.68518252 | 1.27852659 | Eukaryotic tr | ELF5A-2 | | | cytosol |
| AT4G34110 | 5.81377605 | 3.97970503 | 5.83236243 | 4.4916236 | poly(A) bindi | PAB2 | | | nucleus |
| AT1G04985 | 5.74920845 | 3.80514169 | 1.91624882 | 1.33839521 | triacylglycer | | | | mitochondrion |
| AT4G09730 | 5.71763489 | 3.74978744 | 5.76125544 | 4.2699018 | RH39 Chr4:6 | RH39 | | | plastid |
| AT1G29350 | 5.71620072 | 3.73620066 | 3.89824442 | 3.00966257 | RNA polyme | | | | nucleus |
| AT2G20060 | 5.56645992 | 3.38984042 | 3.69788224 | 2.53033914 | Ribosomal p | | | uL4m | mitochondrion |
| AT1G02080 | 5.50318652 | 3.29826196 | 5.53210894 | 3.70179776 | transcription | | | | golgi |
| AT1G80410 | 5.42354471 | 3.16233266 | 5.44637007 | 3.49425752 | tetratricopep | EMB2753 | | | cytosol |
| AT3G44110 | 5.33853966 | 2.96141997 | 2.67574702 | 1.54112823 | DNAJ homol | ATJ3 | | | nucleus |
| AT3G50370 | 5.30874437 | 2.93512327 | 5.35451401 | 3.28256588 | hypothetical | | | | nucleus |
| ATMG00080 | 5.29637704 | 2.722087 | 5.33618216 | 3.0722058 | ribosomal pr | RPL16 | | uL16m | mitochondrion |
| AT5G67500 | 5.21347682 | 2.72623481 | 5.25422254 | 3.05850191 | voltage depe | VDAC2 | | | mitochondrion |
| AT1G69250 | 5.20787129 | 2.54577014 | 5.23121901 | 2.84659499 | Nuclear trans | | | | nucleus |
| AT4G31210 | 5.13834693 | 2.60937201 | 5.14936734 | 2.86303911 | DNA topoiso | | | | plastid |
| AT2G42710 | 5.12133594 | 2.61476742 | 5.15470439 | 2.91249907 | Ribosomal p | | | uL1m | mitochondrion |
| AT3G60240 | 5.12133594 | 2.61476742 | 5.15470439 | 2.91249907 | eukaryotic tr | EIF4G | | | |
| ATMG00560 | 5.11652906 | 2.59414894 | 5.15061387 | 2.86957211 | Nucleic acid- | RPL2 | | | mitochondrion |
| AT3G10690 | 5.04110976 | 5.91721463 | 3.09322417 | 3.281539 | DNA GYRASE | GYRA | | | plastid |
| AT5G60790 | 5.0189646 | 2.47708391 | 5.04439412 | 2.72080512 | ABC transpo | ABCF1 | | | cytosol |
| AT1G68680 | 5.0189646 | 2.47708391 | 3.18913182 | 1.61269051 | SH3/FCH dor | | | | mitochondrion |
| AT1G20010 | 5.00116012 | 2.4170157 | 3.19254711 | 1.5894507 | tubulin beta- | TUBB5 | | | cytosol |
| AT1G54270 | 4.96987911 | 2.34061592 | 3.14223497 | 1.49866458 | eif4a-2 Chr1: | TIF4A-2 | | | cytosol |
| AT4G30930 | 4.90574708 | 2.28251515 | 4.92221257 | 2.49330292 | Ribosomal p | RPL21M | | bL21m | mitochondrion |
| AT3G17465 | 4.90574708 | 2.28251515 | 4.92221257 | 2.49330292 | ribosomal pr | RPL3B | | uL3m | mitochondrion |
| AT3G09820 | 4.90288121 | 2.20423778 | 3.0821281 | 1.42351239 | adenosine ki | ADK1 | | | cytosol |
| AT5G58290 | 4.88796529 | 2.24180612 | 3.06313405 | 1.44142194 | regulatory pa | RPT3 | | | cytosol,nucleus |
| AT2G20140 | 4.84352519 | 2.10692055 | 4.90367498 | 2.35021572 | AAA-type AT | RPT2B | | | cytosol,nucleus |
| AT5G36950 | 4.76885039 | 2.10020504 | 4.78934143 | 2.27301769 | DegP proteas | DEGP10 | | | mitochondrion |
| AT1G48650 | 4.76885039 | 2.10020504 | 4.78934143 | 2.27301769 | DEA(D/H)-bo | | | | mitochondrion,nucleus |
| AT1G31817 | 4.7594055 | 1.91673785 | 4.75323736 | 2.03972893 | Ribosomal L1 | NFD3 | | uS11m | mitochondrion |
| AT5G19820 | 4.75164342 | 2.07412709 | 4.78709483 | 2.2639471 | ARM repeat | emb2734 | | | golgi |
| AT3G13160 | 4.75164342 | 2.07412709 | 4.78709483 | 2.2639471 | Tetratricope | | | mS79 (rPPR3 | mitochondrion |
| AT4G01850 | 4.71835683 | 1.94785252 | 4.77150938 | 2.15665489 | S-adenosylm | SAM2 | | | cytosol,nucleus |
| AT5G19770 | 4.61868299 | 1.90462775 | 2.79151513 | 1.11458662 | tubulin alpha | TUBA3 | | | cytosol |
| AT4G25890 | 4.61355421 | 1.82852378 | 4.62479869 | 1.93615326 | 60S acidic rib | RPP3A | AT4G25890 | | cytosol |
| AT3G18190 | 4.59959938 | 1.88510982 | 2.76379268 | 1.08582445 | TCP-1/cpn60 | CCT4 | | | cytosol |
| AT5G18230 | 4.59959938 | 1.88510982 | 4.64115013 | 2.03486683 | transcription | | | | nucleus |
| AT5G14040 | 4.34952613 | 4.90308999 | 2.27384723 | 1.41687528 | phosphate tr | MPT3 | | | mitochondrion |
| AT3G22330 | 4.28034356 | 6.74472749 | 1.57735231 | 3.11944227 | putative mit | RH53 | | | mitochondrion |
| AT3G13460 | 4.21586129 | 3.40865486 | 6.13003556 | 5.18108282 | evolutionaril | ECT2 | | | cytosol |
| AT2G19750 | 3.49404247 | 1.90462775 | 5.33800707 | 3.10301516 | Ribosomal p | RPS30A | eS30 | | cytosol,nucleus |
| AT3G11130 | 3.44053771 | 2.61190573 | 1.86637846 | 1.20733969 | Clathrin, hea | CHC1 | | | plasma membrane |
| AT2G37230 | 3.36753921 | 2.4898569 | 2.43991258 | 1.50574624 | Tetratricope | | | mL102 (rPPR5) | |
| AT3G23390 | 3.29595512 | 1.60812855 | 5.24044227 | 2.91386851 | Zinc-binding | RPL36AB | eL42 | | cytosol |
| AT2G20580 | 3.00138043 | 1.322121 | 3.07888343 | 1.46501501 | 26S proteaso | RPN1A | | | cytosol,nucleus |
| AT5G17920 | 2.85334897 | 1.13243236 | 2.95180852 | 1.27977251 | Cobalamin-in | MS1 | | | cytosol |
| AT1G75940 | 2.8522268 | 1.7444561 | 2.95441821 | 1.21308738 | Glycosyl hyd | BGLU20 | | | extracellular |
| AT4G36680 | 2.8488001 | 1.82707456 | 1.48981996 | 1.12433316 | Tetratricope | | | mL103 (rPPR | mitochondrion |
| AT5G27850 | 2.78129993 | 1.11127847 | 5.67642348 | 3.85018466 | Ribosomal p | RPL18C | eL18 | | cytosol |
| AT5G53070 | 2.7693092 | 1.06323779 | 4.64115013 | 2.03486683 | Ribosomal p | | | bL9m | mitochondrion |
| AT5G20490 | 2.75530011 | 1.41996775 | 2.77169109 | 1.50103062 | myosin fami | XI-K | | | golgi |
| AT3G52930 | 2.7262609 | 1.70980433 | 2.21112945 | 1.44142194 | Aldolase sup | FBA8 | | | cytosol |
| AT5G62690 | 2.66617561 | 2.53221866 | 1.78677124 | 3.00670145 | tubulin beta | TUBB3 | | | cytosol |
| AT1G48630 | 2.32286869 | 1.13056466 | 6.12031443 | 4.92993135 | receptor for | RACK1B | RACK1 | | cytosol |

**Table S2**: Enriched proteins in FMT co-immunoprecipitations from inflorescences mitochondrial extracts. Statistical analyses based on specific spectral counts identified 75 proteins enriched in IP from mitochondria of at least one FMT/GFP line compared to Col0 IPs (cuts-off FC> 2 and adj-p<0.1)

| AGI | logFC Mito (FMT-GFP vs Col0) | -logadj-p Mito (FMT-GFP vs Col0) | logFC Mito (GFP-FMT vs Col0) | -logadj-p Mito (GFP-FMT vs Col0) | Description | Gene names | SUBA4 | cytoRib (Salih 2020) |
|---|---|---|---|---|---|---|---|---|
| **AT3G52140** | **10.6163518** | **18.7006565** | **11.0379304** | **15.9133601** | **tetratricopep** | **FMT (bait)** | **cytosol** | |
| AT1G43170 | 7.04086998 | 4.53392073 | 6.85698226 | 3.081655 | ribosomal pr | ARP1 | cytosol | uL3 |
| AT1G33120 | 6.50699103 | 3.41240724 | 6.47913514 | 2.67855778 | Ribosomal p | RPL9B | cytosol | uL6 |
| AT5G02870 | 5.89550605 | 2.31420705 | 6.40278116 | 2.67855778 | Ribosomal p | RPL4D | cytosol | uL4 |
| AT4G33010 | 2.75755893 | 0.34819739 | 6.35292694 | 2.60852026 | glycine decar | GLDP1 | mitochondrion | |
| AT1G11860 | #N/A | #N/A | 6.30213954 | 2.50064777 | Glycine cleav | GDCST | mitochondrion | |
| AT5G42080 | 5.61954897 | 1.78820893 | 6.12575342 | 2.6547243 | dynamin-like | DRP1A | plasma membrane | |
| AT1G78570 | 4.19268399 | 0.59256035 | 6.09683909 | 2.6547243 | rhamnose bi | RHM1 | cytosol | |
| AT3G09200 | 5.93142591 | 2.3280676 | 6.06460544 | 2.6547243 | Ribosomal p | RPP0B | cytosol | uL10 |
| AT2G17360 | 6.14628082 | 2.71264881 | 6.06329438 | 2.6547243 | Ribosomal p | RPS4A | cytosol | eS4 |
| AT3G02090 | 3.83505231 | 0.44150636 | 5.89901285 | 2.60852026 | Insulinase (P | At3g02090 | mitochondrion | |
| AT3G09630 | 5.45333534 | 1.53516867 | 5.61729073 | 2.40964479 | Ribosomal p | RPL4A | cytosol | uL4 |
| AT2G36880 | 3.83505231 | 0.44150636 | 5.61508089 | 2.40964479 | methionine a | METK3 | cytosol | |
| AT2G37270 | 4.10602845 | 0.54314488 | 5.57260701 | 2.38524714 | ribosomal pr | RPS5A | cytosol | uS7 |
| AT4G13930 | 4.43427064 | 0.72675827 | 5.52653748 | 2.37568598 | serine hydro | SHM4 | cytosol | |
| AT1G26630 | 3.89121082 | 0.46147239 | 5.43238256 | 2.30258827 | Eukaryotic tr | ELF5A-2 | cytosol | |
| AT2G21870 | #N/A | #N/A | 5.42178632 | 2.13663356 | copper ion bi | At2g21870 | mitochondrion | |
| AT2G34480 | 5.25442223 | 1.378455 | 5.28060433 | 2.15182321 | Ribosomal p | RPL18AB | cytosol | eL20 |
| AT2G20420 | #N/A | #N/A | 5.26535775 | 2.00062032 | ATP citrate ly | At2g20420 | mitochondrion | |
| AT3G48930 | 4.68174102 | 0.90562818 | 5.22777206 | 2.14931911 | Nucleic acid- | RPS11A | cytosol | uS17 |
| AT2G39460 | 4.81438967 | 1.00770051 | 5.22377901 | 2.14931911 | ribosomal pr | RPL23AA | cytosol | uL23 |
| AT1G18080 | 5.1884903 | 1.29925955 | 5.04776569 | 2.00062032 | Transducin/V | RACK1A | cytosol | RACK1 |
| AT3G17390 | 4.81969072 | 1.00770051 | 5.04298181 | 1.99114038 | S-adenosylm | METK4 | plasma membrane,nucleus | |
| AT5G67500 | 3.76679582 | 0.42496618 | 4.99147001 | 1.95215437 | voltage depe | VDAC2 | mitochondrion | |
| AT2G19730 | 4.37004812 | 0.69019113 | 4.98064998 | 1.92819 | Ribosomal L | RPL28A | cytosol | eL28 |
| AT3G53890 | 4.91215179 | 1.06314072 | 4.97529094 | 1.92819 | Ribosomal p | RPS21B | cytosol | eS21 |
| AT2G33210 | #N/A | #N/A | 4.95862187 | 1.84698955 | heat shock p | At2g33210 | mitochondrion | |
| AT3G49010 | 5.34615923 | 1.42055518 | 4.91569747 | 1.88458145 | breast basic | RPL13B | cytosol | eL13 |
| AT4G01100 | 2.75755893 | 0.34819739 | 4.91568598 | 1.88458145 | adenine nucl | ADNT1 | mitochondrion | |
| AT5G65720 | 3.09874377 | 0.34819739 | 4.8411672 | 1.84698955 | nitrogen fixa | NIFS1 | mitochondrion | |
| AT5G15200 | 5.17304711 | 1.29925955 | 4.84114743 | 1.84698955 | Ribosomal p | RPS9B | cytosol | uS4 |
| AT3G18780 | 3.1618086 | 0.34819739 | 4.83362118 | 1.84698955 | actin 2, Sym | ACT2 | cytosol | |
| AT1G14320 | 4.29855076 | 0.65371961 | 4.83359979 | 1.84698955 | Ribosomal p | RPL10A | cytosol | uL16 |
| AT1G08360 | 3.70691262 | 0.40950209 | 4.83357843 | 1.84698955 | Ribosomal p | RPL10AA | cytosol | uL1 |
| AT5G50850 | 2.29478361 | 0.32742234 | 4.80354576 | 1.84698955 | Transketolas | PDH2 | mitochondrion | |
| AT5G27850 | 5.0246756 | 1.18299526 | 4.77066217 | 1.80855389 | Ribosomal p | RPL18C | cytosol | eL18 |
| AT1G41880 | 4.53932604 | 0.80354437 | 4.76190677 | 1.80681212 | Ribosomal p | RPL35AB | cytosol | eL33 |
| AT1G04480 | 4.76998366 | 0.97998029 | 4.75292845 | 1.73087491 | Ribosomal .. | RPL23A | nucleus,cytos | uL14 |
| AT3G23990 | 1.58022103 | 0.32742234 | 4.73553923 | 1.80681212 | heat shock p | CPN60 | mitochondrion | |
| AT5G20290 | 5.04218026 | 1.18299526 | 4.68766753 | 1.73087491 | Ribosomal p | RPS8A | nucleus,cytos | eS8 |
| AT5G48880 | 2.75755893 | 0.34819739 | 4.67724435 | 1.71331221 | peroxisomal | KAT5 | peroxisome | |
| AT2G18020 | 4.52448975 | 0.77335686 | 4.67722291 | 1.71331221 | Ribosomal p | RPL8A | cytosol | uL2 |
| AT2G05710 | #N/A | #N/A | 4.66564603 | 1.71331221 | aconitase 3, | ACO2 | mitochondrion | |
| AT5G66760 | #N/A | #N/A | 4.66564603 | 1.71331221 | succinate de | SDH1-1 | mitochondrion | |
| AT2G41840 | 5.22181367 | 1.29925955 | 4.58876034 | 1.57688947 | Ribosomal p | RPS2C | cytosol | uS5 |
| AT2G27530 | 4.63396316 | 0.87222052 | 4.58876034 | 1.57688947 | Ribosomal p | RPL10AB | cytosol | uL1 |
| AT3G52300 | #N/A | #N/A | 4.58214066 | 1.60524477 | ATP synthase | At3g52300 | mitochondrion | |
| AT1G48030 | #N/A | #N/A | 4.58214066 | 1.60524477 | mitochondria | LPD1 | mitochondrion | |
| AT5G37510 | #N/A | #N/A | 4.58214066 | 1.60524477 | NADH-ubiqui | EMB1467 | mitochondrion | |
| AT5G27700 | 4.10602845 | 0.54314488 | 4.504392 | 1.45790745 | Ribosomal p | RPS21C | cytosol | eS21 |
| AT1G04985 | 3.76679582 | 0.42496618 | 4.504392 | 1.45790745 | Triacylglycer | At1g04985 F | mitochondrion | |
| AT3G04840 | 3.59076925 | 0.39795412 | 4.504392 | 1.45790745 | Ribosomal p | RPS3AA | cytosol | eS1 |
| AT1G10290 | 3.42839881 | 0.38653049 | 4.504392 | 1.45790745 | dynamin-like | DRP2A | golgi | |
| AT1G24020 | 2.43850297 | 0.32742234 | 4.504392 | 1.45790745 | MLP-like pro | MLP423 | golgi | |
| AT1G72370 | 4.36148445 | 2.71264881 | 4.49180229 | 2.32970023 | 40s riboso | RPSaA | cytosol | uS2 |
| AT4G31700 | 4.68176815 | 0.90562818 | 4.4169572 | 1.34059863 | ribosomal pr | RPS6A | cytosol | eS6 |
| AT1G02780 | 4.68132506 | 0.89290971 | 4.4169572 | 1.34059863 | Ribosomal p | RPL19A | cytosol | eL19 |
| AT5G52840 | #N/A | #N/A | 4.398229 | 1.34676601 | NADH-ubiqui | At5g52840 | mitochondrion | |
| AT5G10860 | #N/A | #N/A | 4.38147294 | 1.34059863 | Cystathionine | CBSX3 | mitochondrion | |
| AT2G45710 | 5.32605844 | 1.42055518 | 4.30706456 | 1.16088388 | Zinc-binding | RPS27A | cytosol | eS27 |
| AT5G10360 | 5.01280181 | 1.18299526 | 4.30706456 | 1.16088388 | Ribosomal p | RPS6B | cytosol | eS6 |
| AT2G07698 | 2.27541202 | 0.40920168 | 4.29144092 | 2.14931911 | ATPase, F1 c | | plastid,mitochondrion | |
| AT4G34670 | 3.94432948 | 0.46147239 | 4.23292919 | 1.07490384 | Ribosomal p | RPS3AB | cytosol | eS1 |
| AT1G56070 | 2.64557597 | 1.03128762 | 4.11636447 | 2.3844326 | Ribosomal p | LOS1 | cytosol | |
| AT3G22330 | 3.56297934 | 1.42055518 | 4.02211963 | 1.92819 | putative mito | RH53 | mitochondrion | |
| AT1G22780 | 3.69738332 | 1.53516867 | 3.93102504 | 1.87296993 | Ribosomal .. | RPS18A | cytosol | uS13 |
| AT3G45030 | 5.03721911 | 1.18299526 | 3.64206432 | 0.45996701 | Ribosomal p | RPS20A | cytosol | uS10 |
| AT4G37930 | 0.61330629 | 0.15015542 | 3.50816319 | 1.83574926 | serine transh | SHM1 | mitochondrion | |
| AT2G42740 | 5.11097946 | 1.21080283 | 3.28210613 | 0.38543793 | ribosomal pr | RPL11A | cytosol | uL5 |
| AT1G04820 | 2.55750403 | 0.49894327 | 3.26366067 | 1.45790745 | tubulin alpha | | cytosol | |
| AT5G08670 | 0.37408977 | 0.10412212 | 3.1675491 | 1.67794684 | ATP synthase | At5g08670 | mitochondrion | |
| AT5G62690 | 2.87215926 | 1.22017381 | 2.93858013 | 1.33645031 | tubulin beta | | cytosol | |
| AT5G46800 | 2.27972926 | 1.15289727 | 2.84167279 | 1.47005683 | Mitochondria | BOU | mitochondrion | |
| AT4G02930 | 1.88758392 | 0.42096367 | 2.71630852 | 1.16088388 | GTP binding | TUFA | mitochondrion | |
| AT1G07920 | 1.64750467 | 0.66383692 | 2.61335852 | 1.43164769 | | | cytosol | |
| AT1G20620 | 2.57550381 | 1.49802188 | 2.38125367 | 1.09806956 | catalase 3, S | CAT3 | peroxisome | |

**Table S3:** Enriched proteins in FMT co-immunoprecipitation from seedling mitochondrial extracts. One hundred and twenty-four proteins were enriched in GFP-FMT compared to MS2-GFP IPs, with at least 3 spectra and a FC above 2, among them 51 were cytosolic ribosomal proteins

| Accession | Description | SUBA4 localization | cyto Rib, Salih2020 | Specific spectra_ Col0 | Specific spectra_ GFP-FMT | Log2FC | Z-score Col0 | Z-score GFP-FMT | Uniprot Accession |
|---|---|---|---|---|---|---|---|---|---|
| AT1G08360 | Ribosomal p | cytosol | **L10A** | 0 | 3 | 2.3971 | -0.2643 | -0.0070 | R10A1_ARATH |
| AT3G53430 | Ribosomal p | cytosol | **L12** | 0 | 4 | 2.7976 | -0.2458 | 0.5215 | RL122_ARATH |
| AT3G49010 | breast basic | cytosol | **L13** | 0 | 7 | 3.5861 | -0.2613 | 1.0173 | RL131_ARATH |
| AT3G24830 | Ribosomal p | cytosol | **L13A** | 0 | 3 | 2.3971 | -0.2613 | 0.0287 | R13A2_ARATH |
| AT4G27090 | Ribosomal p | cytosol | **L14** | 0 | 4 | 2.7976 | -0.2268 | 0.8236 | RL142_ARATH |
| AT4G16720 | Ribosomal p | cytosol | **L15** | 0 | 7 | 3.5861 | -0.2606 | 1.0346 | RL151_ARATH |
| AT1G27400 | Ribosomal p | plastid,nucle | **L17** | 0 | 6 | 3.3680 | -0.2503 | 1.0282 | RL171_ARATH |
| AT3G05590 | ribosomal pr | cytosol | **L18** | 0 | 3 | 2.3971 | -0.2548 | 0.1072 | RL182_ARATH |
| AT2G34480 | Ribosomal p | cytosol | **L18A** | 0 | 12 | 4.3532 | -0.2512 | 2.7244 | R18A2_ARATH |
| AT1G04480 | Ribosomal .. | nucleus,cytos | **L23** | 0 | 5 | 3.1108 | -0.2310 | 1.1201 | RL23_ARATH |
| AT2G39460 | ribosomal pr | cytosol | **L23A** | 0 | 13 | 4.4675 | -0.2396 | 3.5955 | R23A1_ARATH |
| AT2G36620 | ribosomal pr | cytosol | **L24** | 0 | 8 | 3.7756 | -0.2448 | 1.7787 | RL241_ARATH |
| AT3G49910 | Translation p | cytosol | **L26** | 0 | 6 | 3.3680 | -0.2349 | 1.3923 | RL261_ARATH |
| AT1G70600 | Ribosomal p | cytosol | **L27A** | 0 | 4 | 2.7976 | -0.2349 | 0.6948 | R27A3_ARATH |
| AT2G19730 | Ribosomal L | cytosol | **L28** | 0 | 6 | 3.3680 | -0.2330 | 1.4371 | RL281_ARATH |
| AT1G43170 | ribosomal pr | cytosol | **L3** | 0 | 20 | 5.0842 | -0.2915 | 1.8903 | RL31_ARATH |
| AT1G36240 | Ribosomal p | cytosol | **L30** | 0 | 4 | 2.7976 | -0.2074 | 1.1315 | RL301_ARATH |
| AT4G18100 | Ribosomal p | cytosol | **L32** | 0 | 9 | 3.9431 | -0.2261 | 2.7494 | RL321_ARATH |
| AT1G26880 | Ribosomal p | nucleus,cytos | **L34** | 0 | 3 | 2.3971 | -0.2153 | 0.5822 | RL341_ARATH |
| AT3G53740 | Ribosomal p | cytosol | **L36** | 0 | 5 | 3.1108 | -0.1971 | 1.7896 | RL362_ARATH |
| AT3G60245 | Zinc-binding | cytosol | **L37A** | 0 | 3 | 2.3971 | -0.1818 | 0.9857 | R37A2_ARATH |
| AT3G09630 | Ribosomal p | cytosol | **L4** | 0 | 9 | 3.9431 | -0.2929 | 0.4007 | RL4A_ARATH |
| AT5G02870 | Ribosomal p | cytosol | **L4** | 0 | 9 | 3.9431 | -0.2930 | 0.3979 | RL4B_ARATH |
| AT3G25520 | ribosomal pr | cytosol | **L5** | 0 | 11 | 4.2290 | -0.2815 | 1.1381 | RL51_ARATH |
| AT1G74050 | Ribosomal p | cytosol | **L6** | 0 | 5 | 3.1108 | -0.2687 | 0.3762 | RL63_ARATH |
| AT2G44120 | Ribosomal p | cytosol | **L7** | 0 | 7 | 3.5861 | -0.2708 | 0.7554 | RL73_ARATH |
| AT2G47610 | Ribosomal p | cytosol | **L7A** | 0 | 7 | 3.5861 | -0.2740 | 0.6679 | RL7A1_ARATH |
| AT2G18020 | Ribosomal p | cytosol | **L8** | 0 | 7 | 3.5861 | -0.2742 | 0.6624 | RL81_ARATH |
| AT1G33120 | Ribosomal p | cytosol | **L9** | 0 | 13 | 4.4675 | -0.2573 | 2.7008 | RL91_ARATH |
| AT3G09200 | Ribosomal p | cytosol | **P0** | 0 | 9 | 3.9431 | -0.2841 | 0.7082 | RLA02_ARATH |
| AT2G27710 | 60S acidic rib | cytosol | **P2** | 0 | 7 | 3.5861 | -0.2105 | 2.4107 | RLA22_ARATH |
| AT1G18080 | Transducin/V | cytosol | **RACK1** | 0 | 8 | 3.7756 | -0.2850 | 0.5214 | GBLPA_ARATH |
| AT4G25740 | RNA binding | cytosol | **S10** | 0 | 5 | 3.1108 | -0.2508 | 0.7305 | RS101_ARATH |
| AT3G48930 | Nucleic acid- | cytosol | **S11** | 0 | 12 | 4.3532 | -0.2428 | 3.1146 | RS111_ARATH |
| AT1G04270 | cytosolic ribo | cytosol | **S15** | 0 | 4 | 2.7976 | -0.2385 | 0.6381 | RS151_ARATH |
| AT5G18380 | Ribosomal p | cytosol | **S16** | 0 | 5 | 3.1108 | -0.2349 | 1.0435 | RS163_ARATH |
| AT1G22780 | Ribosomal .. | cytosol | **S18** | 0 | 12 | 4.3532 | -0.2385 | 3.3176 | RS18_ARATH |
| AT1G58380 | Ribosomal p | cytosol | **S2** | 0 | 5 | 3.1108 | -0.2789 | 0.1751 | RS21_ARATH |
| AT2G41840 | Ribosomal p | cytosol | **S2** | 0 | 4 | 2.7976 | -0.2791 | -0.0067 | RS23_ARATH |
| AT3G45030 | Ribosomal p | cytosol | **S20** | 0 | 9 | 3.9431 | -0.2188 | 3.0029 | RS201_ARATH |
| AT3G09680 | Ribosomal p | cytosol | **S23** | 0 | 9 | 3.9431 | -0.2324 | 2.5280 | |
| AT2G40510 | Ribosomal p | cytosol | **S26** | 0 | 7 | 3.5861 | -0.2261 | 1.9838 | RS262_ARATH |
| AT2G45710 | Zinc-binding | cytosol | **S27** | 0 | 4 | 2.7976 | -0.1681 | 1.7565 | RS271_ARATH |
| AT3G04840 | Ribosomal p | cytosol | **S3A** | 0 | 10 | 4.0931 | -0.2750 | 1.2239 | RS3A1_ARATH |
| AT4G34670 | Ribosomal p | cytosol | **S3A** | 0 | 4 | 2.7976 | -0.2750 | 0.0580 | RS3A2_ARATH |
| AT2G17360 | Ribosomal p | cytosol | **S4** | 0 | 9 | 3.9431 | -0.2748 | 1.0364 | RS41_ARATH |
| AT2G37270 | ribosomal pr | cytosol | **S5** | 0 | 3 | 2.3971 | -0.2616 | 0.0250 | RS51_ARATH |
| AT5G10360 | Ribosomal p | cytosol | **S6** | 0 | 3 | 2.3971 | -0.2724 | -0.1046 | RS62_ARATH |
| AT5G20290 | Ribosomal p | nucleus,cytos | **S8** | 0 | 8 | 3.7756 | -0.2659 | 1.1197 | RS81_ARATH |
| AT5G15200 | Ribosomal p | cytosol | **S9** | 0 | 4 | 2.7976 | -0.2587 | 0.3171 | RS91_ARATH |
| AT1G72370 | 40s ribosoma | cytosol | **SA** | 0 | 19 | 5.0106 | -0.2811 | 2.5238 | RSSA1_ARATH |
| AT3G52140 | tetratricopep | cytosol | | 0 | 209 | 8.4615 | -0.3160 | 6.8451 | FMT (bait) |
| AT1G54270 | eif4a-2, Sym | cytosol | | 1 | 25 | 2.2344 | -0.0367 | 2.3612 | IF4A2_ARATH |
| AT2G36880 | methionine a | cytosol | | 0 | 19 | 5.0106 | -0.2916 | 1.7530 | METK3_ARATH |
| AT1G56070 | Ribosomal p | cytosol | | 1 | 29 | 2.4475 | -0.1843 | 1.0154 | EF2_ARATH |
| AT5G17920 | Cobalamin-in | cytosol | | 0 | 25 | 5.4043 | -0.3082 | 0.9285 | METE1_ARATH |
| AT5G19550 | aspartate am | cytosol | | 0 | 13 | 4.4675 | -0.2928 | 0.9063 | AAT2_ARATH |
| AT5G44020 | HAD superfa | cytosol | | 0 | 7 | 3.5861 | -0.2769 | 0.5900 | Q9FNC4_ARATH |
| AT1G26630 | Eukaryotic tr | cytosol | | 0 | 4 | 2.7976 | -0.2423 | 0.5772 | IF5A2_ARATH |
| AT3G16460 | Mannose-bin | cytosol | | 1 | 18 | 1.7632 | -0.1567 | 0.5653 | JAL34_ARATH |
| AT2G30860 | glutathione S | cytosol | | 0 | 5 | 3.1108 | -0.2640 | 0.4700 | GSTF9_ARATH |
| AT5G54160 | O-methyltran | cytosol | | 0 | 8 | 3.7756 | -0.2890 | 0.3960 | OMT1_ARATH |
| AT4G09000 | general regu | cytosol | | 0 | 5 | 3.1108 | -0.2759 | 0.2336 | 14331_ARATH |

| Accession | Description | SUBA4 localization | cyto Rib, Salih2020 | Specific spectra_ Col0 | Specific spectra_ GFP-FMT | Log2FC | Z-score Col0 | Z-score GFP-FMT | Uniprot Accession |
|---|---|---|---|---|---|---|---|---|---|
| AT4G13930 | serine hydrox | cytosol | | 0 | 8 | 3.7756 | -0.2974 | 0.1347 | GLYC4_ARATH |
| AT1G52100 | Mannose-bin | cytosol | | 0 | 7 | 3.5861 | -0.2952 | 0.0865 | JAL11_ARATH |
| AT5G62690 | tubulin beta | cytosol | | 0 | 6 | 3.3680 | -0.2961 | -0.0506 | |
| AT5G56010 | heat shock p | cytosol | | 0 | 9 | 3.9431 | -0.3065 | -0.0790 | HS903_ARATH |
| AT5G10450 | G-box regula | cytosol | | 0 | 3 | 2.3971 | -0.2721 | -0.1021 | 14336_ARATH |
| AT5G66420 | unknown pro | cytosol | | 0 | 7 | 3.5861 | -0.3053 | -0.1898 | Q9FJZ7_ARATH |
| AT2G36530 | Enolase, Sym | cytosol | | 0 | 4 | 2.7976 | -0.2957 | -0.2706 | ENO2_ARATH |
| AT3G16410 | nitrile specif | cytosol | | 0 | 5 | 3.1108 | -0.3041 | -0.3221 | JAL29_ARATH |
| AT1G02500 | S-adenosylm | cytosol | | 0 | 3 | 2.3971 | -0.2918 | -0.3388 | METK1_ARATH |
| AT1G79550 | phosphoglyce | cytosol | | 0 | 3 | 2.3971 | -0.2925 | -0.3468 | PGKY3_ARATH |
| AT5G36230 | ARM repeat | cytosol | | 0 | 3 | 2.3971 | -0.2933 | -0.3565 | Q93ZC2_ARATH |
| AT1G57720 | Translation e | cytosol | | 0 | 3 | 2.3971 | -0.2934 | -0.3584 | EF1G2_ARATH |
| AT1G09640 | Translation e | cytosol | | 0 | 3 | 2.3971 | -0.2935 | -0.3593 | EF1G1_ARATH |
| AT4G31480 | Coatomer, be | cytosol | | 0 | 7 | 3.5861 | -0.3115 | -0.3610 | |
| AT3G55610 | delta 1-pyrro | cytosol | | 0 | 5 | 3.1108 | -0.3072 | -0.3842 | P5CS2_ARATH |
| AT5G02500 | heat shock co | cytosol | | 0 | 4 | 2.7976 | -0.3051 | -0.4210 | MD37E_ARATH |
| AT1G16020 | Protein of un | cytosol | | 0 | 3 | 2.3971 | -0.2998 | -0.4347 | |
| AT3G15980 | Coatomer, be | cytosol | | 0 | 4 | 2.7976 | -0.3109 | -0.5126 | COB23_ARATH |
| AT2G21390 | Coatomer, al | cytosol | | 0 | 3 | 2.3971 | -0.3146 | -0.6130 | COPA2_ARATH |
| AT1G21750 | PDI-like 1-1, | endoplasmic reticulum | | 0 | 8 | 3.7756 | -0.2991 | 0.0821 | PDI11_ARATH |
| AT1G66280 | Glycosyl hydr | endoplasmic reticulum | | 0 | 7 | 3.5861 | -0.3002 | -0.0513 | BGL22_ARATH |
| AT1G66270 | Glycosyl hydr | endoplasmic reticulum | | 0 | 4 | 2.7976 | -0.3002 | -0.3428 | BGL21_ARATH |
| AT5G28540 | heat shock p | endoplasmic reticulum | | 0 | 3 | 2.3971 | -0.3057 | -0.5058 | MD37A_ARATH |
| AT4G27150 | seed storage | extracellular | | 0 | 3 | 2.3971 | -0.2477 | 0.1923 | 2SS2_ARATH |
| AT5G26260 | TRAF-like fa | extracellular | | 0 | 4 | 2.7976 | -0.2878 | -0.1453 | O81493_ARATH |
| AT1G30630 | Coatomer ep | golgi | | 0 | 4 | 2.7976 | -0.2802 | -0.0244 | COPE1_ARATH |
| AT1G10290 | dynamin-like | golgi | | 0 | 4 | 2.7976 | -0.3110 | -0.5138 | DRP2A_ARATH |
| AT5G20490 | Myosin fami | golgi | | 0 | 3 | 2.3971 | -0.3169 | -0.6406 | Q3E994_ARATH |
| AT3G01280 | voltage depe | mitochondrion | | 0 | 7 | 3.5861 | -0.2776 | 0.5707 | VDAC1_ARATH |
| AT4G02930 | GTP binding | mitochondrion | | 1 | 11 | 1.0591 | -0.0634 | 0.5040 | EFTM_ARATH |
| AT4G01100 | adenine nucl | mitochondrion | | 0 | 7 | 3.5861 | -0.2879 | 0.2869 | ADNT1_ARATH |
| AT3G48680 | gamma carb | mitochondrion | | 0 | 3 | 2.3971 | -0.2738 | -0.1221 | GCAL2_ARATH |
| AT3G54110 | plant uncoup | mitochondrion | | 0 | 3 | 2.3971 | -0.2823 | -0.2237 | PUMP1_ARATH |
| AT5G40810 | Cytochrome ( | mitochondrion | | 0 | 3 | 2.3971 | -0.2824 | -0.2254 | CYC1B_ARATH |
| AT4G29130 | hexokinase 1 | mitochondrion | | 0 | 3 | 2.3971 | -0.2988 | -0.4228 | HXK1_ARATH |
| AT3G22330 | putative mito | mitochondrion | | 0 | 3 | 2.3971 | -0.3040 | -0.4853 | RH53_ARATH |
| AT4G39260 | cold, circadia | nucleus | | 0 | 3 | 2.3971 | -0.2472 | 0.1978 | RBG8_ARATH |
| AT5G07350 | TUDOR-SN p | nucleus | | 0 | 3 | 2.3971 | -0.3121 | -0.5831 | TSN1_ARATH |
| AT3G04120 | glyceraldehy | nucleus,cytosol | | 1 | 12 | 1.1832 | 0.0265 | 1.0827 | G3PC1_ARATH |
| AT4G35090 | catalase 2, S | peroxisome | | 1 | 17 | 1.6814 | -0.0836 | 1.0285 | CATA2_ARATH |
| AT1G06460 | alpha-crystal | peroxisome | | 0 | 7 | 3.5861 | -0.2791 | 0.5292 | Q208N7_ARATH |
| AT1G54340 | isocitrate de | peroxisome | | 0 | 4 | 2.7976 | -0.2937 | -0.2388 | ICDHX_ARATH |
| AT2G33150 | peroxisomal | peroxisome | | 0 | 3 | 2.3971 | -0.2968 | -0.3992 | THIK2_ARATH |
| AT3G06810 | acyl-CoA deh | peroxisome | | 0 | 3 | 2.3971 | -0.3094 | -0.5505 | IBR3_ARATH |
| AT3G11130 | Clathrin, hea | plasma membrane | | 0 | 32 | 5.7589 | -0.3177 | 0.2157 | CLAH1_ARATH |
| AT5G42080 | dynamin-like | plasma membrane | | 0 | 8 | 3.7756 | -0.3038 | -0.0655 | DRP1A_ARATH |
| AT1G59610 | dynamin-like | plasma membrane | | 0 | 8 | 3.7756 | -0.3111 | -0.2940 | DRP2B_ARATH |
| AT2G18960 | H(+)-ATPase | plasma membrane | | 0 | 3 | 2.3971 | -0.3115 | -0.5760 | PMA1_ARATH |
| AT3G17390 | S-adenosylm | plasma membrane,nucle | | 1 | 24 | 2.1758 | -0.0227 | 2.3817 | METK4_ARATH |
| AT2G47730 | glutathione S | plastid | | 0 | 9 | 3.9431 | -0.2752 | 1.0228 | GSTF8_ARATH |
| AT1G67090 | ribulose bisp | plastid | | 0 | 3 | 2.3971 | -0.2520 | 0.1403 | RBS1A_ARATH |
| ATCG00490 | ribulose-bisp | plastid | | 0 | 6 | 3.3680 | -0.2979 | -0.0926 | RBL_ARATH |
| AT2G39730 | rubisco activ | plastid | | 0 | 5 | 3.1108 | -0.2976 | -0.1931 | RCA_ARATH |
| AT3G58610 | ketol-acid re | plastid | | 0 | 3 | 2.3971 | -0.3031 | -0.4744 | ILV5_ARATH |
| AT3G14210 | epithiospecif | vacuole | | 0 | 11 | 4.2290 | -0.2917 | 0.7013 | |
| AT1G54000 | GDSL-like Lip | vacuole | | 0 | 10 | 4.0931 | -0.2916 | 0.5748 | GDL18_ARATH |
| AT1G04040 | HAD superfa | vacuole | | 0 | 6 | 3.3680 | -0.2767 | 0.4071 | Q9ZWC4_ARATH |
| AT1G54010 | GDSL-like Lip | vacuole | | 0 | 5 | 3.1108 | -0.2912 | -0.0676 | GDL19_ARATH |
| AT1G20260 | ATPase, V1 c | vacuole | | 0 | 6 | 3.3680 | -0.2983 | -0.1033 | VATB3_ARATH |
| AT1G04410 | Lactate/mala | vacuole | | 0 | 3 | 2.3971 | -0.2856 | -0.2644 | MDHC1_ARATH |
| AT1G78900 | vacuolar ATP | vacuole,golgi | | 0 | 6 | 3.3680 | -0.3042 | -0.2431 | VATA_ARATH |
| AT3G42050 | vacuolar ATP | vacuole,golgi | | 0 | 3 | 2.3971 | -0.2955 | -0.3828 | VATH_ARATH |

**Table S4:** Enriched proteins in FMT co-immunoprecipitation from seedling total extracts after formaldehyde crosslinking. Statistical analyses based on specific spectral counts identified 119 proteins enriched in IP from crosslinked seedlings total extracts of FMT-GFP line compared to MS2-GFP IPs (cuts-off FC> 2 and adj-p<0.1)

| AGI | logFC CL-TOT (FMT-GFP vs MS2-GFP) | -logadj-p CL-TOT (FMT-GFP vs MS2-GFP) | logFC CL-TOT (GFP-FMT vs MS2-GFP) | -logadj-p CL-TOT (GFP-FMT vs MS2-GFP) | Description | SUBA4 localization | cyto RIB Salih,2020 |
|---|---|---|---|---|---|---|---|
| **AT3G52140** | **7.496** | **55.774** | **7.385** | **6.265** | **FMT (bait)** | **cytosol** | |
| AT1G14320 | 1.757 | 1.391 | | | Ribosomal p | cytosol | L10 |
| AT2G27530 | 4.899 | 3.176 | | | Ribosomal p | cytosol | L10A |
| AT1G08360 | 2.459 | 1.442 | | | Ribosomal p | cytosol | L10A |
| AT2G42740 | 2.308 | 2.693 | | | ribosomal pr | cytosol | L11 |
| AT3G49010 | 2.119 | 1.772 | | | breast basic | cytosol | L13 |
| AT3G07110 | 3.777 | 2.449 | | | Ribosomal p | cytosol | L13A |
| AT3G24830 | 2.913 | 1.817 | | | Ribosomal p | cytosol | L13A |
| AT4G27090 | 3.512 | 2.999 | | | Ribosomal p | cytosol | L14 |
| AT1G67430 | 2.584 | 1.362 | | | Ribosomal p | nucleus,cytos | L17 |
| AT1G27400 | 2.466 | 1.301 | | | Ribosomal p | plastid,nucle | L17 |
| AT5G27850 | 5.102 | 2.306 | | | Ribosomal p | cytosol | L18 |
| AT2G34480 | 2.557 | 2.772 | | | Ribosomal p | cytosol | L18A |
| AT3G16780 | 5.260 | 2.954 | | | Ribosomal p | cytosol | L19 |
| AT4G02230 | 4.878 | 2.375 | | | Ribosomal p | cytosol | L19 |
| AT1G02780 | 2.580 | 2.605 | | | Ribosomal p | cytosol | L19 |
| AT1G04480 | 2.865 | 2.569 | | | Ribosomal .. | nucleus,cytos | L23 |
| AT2G39460 | 2.915 | 3.984 | | | ribosomal pr | cytosol | L23A |
| AT3G49910 | 3.586 | 3.040 | | | Translation p | cytosol | L26 |
| AT4G15000 | 5.054 | 1.959 | | | Ribosomal L | cytosol | L27 |
| AT1G23290 | 5.256 | 2.630 | | | Ribosomal p | cytosol | L27A |
| AT1G70600 | 4.268 | 2.294 | | | Ribosomal p | cytosol | L27A |
| AT4G29410 | 3.284 | 2.194 | | | Ribosomal L | cytosol | L28 |
| AT2G19730 | 2.638 | 3.323 | | | Ribosomal L | cytosol | L28 |
| AT1G43170 | 1.977 | 3.122 | | | ribosomal pr | cytosol | L3 |
| AT4G18100 | 2.411 | 2.000 | | | Ribosomal p | cytosol | L32 |
| AT1G69620 | 4.969 | 1.965 | | | ribosomal pr | cytosol | L34 |
| AT3G09500 | 3.966 | 3.022 | | | Ribosomal L | cytosol | L35 |
| AT3G23390 | 5.000 | 1.530 | | | Zinc-binding | cytosol | L36A |
| AT1G52300 | 2.752 | 1.450 | | | Zinc-binding | cytosol | L37 |
| AT2G43460 | 4.556 | 2.496 | | | Ribosomal L | cytosol | L38 |
| AT3G09630 | 2.336 | 3.542 | | | Ribosomal p | cytosol | L4 |
| AT5G02870 | 2.158 | 2.997 | | | Ribosomal p | cytosol | L4 |
| AT5G39740 | 5.081 | 2.740 | | | ribosomal pr | cytosol | L5 |
| AT3G25520 | 3.241 | 2.887 | | | ribosomal pr | cytosol | L5 |
| AT1G18540 | 2.201 | 2.141 | | | Ribosomal p | cytosol | L6 |
| AT2G01250 | 2.740 | 3.438 | | | Ribosomal p | cytosol | L7 |
| AT2G44120 | 2.695 | 2.675 | | | Ribosomal p | cytosol | L7 |
| AT3G62870 | 6.092 | 3.150 | | | Ribosomal p | cytosol | L7A |
| AT2G47610 | 3.093 | 1.526 | | | Ribosomal p | cytosol | L7A |
| AT2G18020 | 3.200 | 2.401 | | | Ribosomal p | cytosol | L8 |
| AT4G36130 | 3.165 | 2.152 | | | Ribosomal p | cytosol | L8 |
| AT1G33120 | 2.441 | 4.672 | | | Ribosomal p | cytosol | L9 |
| AT3G09200 | 2.707 | 2.872 | | | Ribosomal p | cytosol | P0 |
| AT2G27710 | 4.792 | 2.293 | | | 60S acidic rib | cytosol | P2 |
| AT1G18080 | 2.188 | 2.630 | | | Transducin/W | cytosol | RACK1 |
| AT5G41520 | 3.795 | 2.569 | | | RNA binding | cytosol | S10 |
| AT5G23740 | 5.732 | 2.799 | | | ribosomal pr | cytosol | S11 |
| AT3G48930 | 2.907 | 3.155 | | | Nucleic acid- | cytosol | S11 |
| AT1G15930 | 2.496 | 1.567 | | | Ribosomal p | cytosol | S12 |
| AT3G60770 | 2.558 | 1.539 | | | Ribosomal p | cytosol | S13 |
| AT3G11510 | 2.829 | 1.338 | | | Ribosomal p | cytosol | S14 |
| AT1G07770 | 2.273 | 2.209 | | | ribosomal pr | cytosol | S15A |
| AT2G04390 | 3.483 | 4.719 | | | Ribosomal S | cytosol | S17 |
| AT1G22780 | 2.086 | 4.472 | | | Ribosomal .. | cytosol | S18 |
| AT3G02080 | 3.342 | 3.141 | | | Ribosomal p | cytosol | S19 |
| AT5G61170 | 2.538 | 1.085 | | | Ribosomal p | cytosol | S19 |
| AT1G58380 | 2.420 | 1.447 | | | Ribosomal p | cytosol | S2 |
| AT2G41840 | 1.916 | 1.499 | | | Ribosomal p | cytosol | S2 |
| AT3G45030 | 1.979 | 1.261 | | | Ribosomal p | cytosol | S20 |
| AT3G53890 | 2.787 | 1.883 | | | Ribosomal p | cytosol | S21 |
| AT5G02960 | 3.272 | 3.147 | | | Ribosomal p | cytosol | S23 |
| AT5G28060 | 3.253 | 2.292 | | | Ribosomal p | cytosol | S24 |
| AT3G56340 | 5.021 | 2.558 | | | Ribosomal p | cytosol | S26 |
| AT3G10090 | 1.897 | 1.396 | | | Nucleic acid- | cytosol | S28 |
| AT3G43980 | 3.259 | 2.310 | | | 0 | nucleus,cytos | S29 |
| AT3G53870 | 4.128 | 2.148 | | | Ribosomal p | cytosol | S3 |
| AT5G35530 | 3.845 | 2.310 | | | Ribosomal p | cytosol | S3 |
| AT4G34670 | 3.568 | 4.179 | | | Ribosomal p | cytosol | S3A |
| AT3G04840 | 2.164 | 2.682 | | | Ribosomal p | cytosol | S3A |
| AT2G17360 | 2.468 | 2.772 | | | Ribosomal p | cytosol | S4 |
| AT2G37270 | 2.466 | 1.902 | | | ribosomal pr | cytosol | S5 |
| AT3G11940 | 2.005 | 1.508 | | | ribosomal pr | cytosol | S5 |
| AT5G10360 | 2.381 | 3.588 | | | Ribosomal p | cytosol | S6 |
| AT4G31700 | 2.337 | 4.146 | | | ribosomal pr | cytosol | S6 |
| AT3G02560 | 5.920 | 3.375 | | | Ribosomal p | cytosol | S7 |
| AT5G16130 | 3.797 | 2.596 | | | Ribosomal p | cytosol | S7 |
| AT1G48830 | 3.451 | 2.230 | | | Ribosomal p | cytosol | S7 |
| AT5G20290 | 3.206 | 4.928 | | | Ribosomal p | nucleus,cytos | S8 |
| AT5G15200 | 2.195 | 2.097 | | | Ribosomal p | cytosol | S9 |
| AT1G72370 | 1.808 | 2.401 | | | 40s ribosoma | cytosol | SA |
| AT3G52200 | 6.752 | 5.305 | 6.028 | 1.169 | Dihydrolipoa | mitochondrion | |
| AT1G54220 | 5.991 | 3.540 | 5.898 | 1.187 | Dihydrolipoa | mitochondrion | |
| AT4G18070 | 5.964 | 2.806 | | | unknown pro | nucleus | |
| AT3G20650 | 5.861 | 2.799 | | | mRNA cappi | nucleus | |
| AT1G31440 | 5.506 | 3.608 | | | SH3 domain- | plasma membrane | |
| AT3G13930 | 5.313 | 14.255 | 4.481 | 1.147 | Dihydrolipoa | mitochondrion | |
| AT3G29410 | 5.300 | 2.310 | | | Terpenoid cy | plastid | |
| AT4G26970 | 5.022 | 2.466 | | | aconitase 2, | mitochondrion | |
| AT2G25670 | 4.991 | 2.373 | | | unknown pro | cytosol | |
| AT1G16350 | 4.930 | 2.455 | | | Aldolase-typ | peroxisome | |
| AT3G18080 | 4.848 | 2.283 | | | B-S glucosida | extracellular | |
| AT1G69250 | 4.632 | 3.727 | | | Nuclear trans | nucleus | |
| AT1G64330 | 4.016 | 1.747 | | | myosin heav | vacuole | |
| AT4G17620 | 3.870 | 1.998 | | | glycine-rich p | cytosol | |
| AT3G12390 | 3.738 | 2.673 | | | Nascent poly | nucleus | |
| AT5G52040 | 3.648 | 1.801 | | | RNA-binding | nucleus | |
| AT1G67680 | 3.634 | 1.478 | | | SRP72 RNA-b | nucleus | |
| AT4G26780 | 3.215 | 1.574 | | | Co-chaperon | mitochondrion | |
| AT2G20280 | 3.003 | 1.238 | | | Zinc finger C | nucleus | |
| AT5G22650 | 2.781 | 1.430 | | | histone deac | nucleus | |
| AT2G29560 | 2.749 | 2.102 | | | cytosolic eno | cytosol | |
| AT4G36700 | 2.715 | 1.049 | | | RmlC-like cu | extracellular | |
| AT1G03890 | 2.693 | 2.295 | 2.457 | | RmlC-like cu | extracellular | |
| AT1G76810 | 2.688 | 1.909 | | | eukaryotic tr | cytosol | |
| AT5G41790 | 2.679 | 1.217 | | | COP1-interac | cytosol | |
| AT1G26550 | 2.594 | 1.395 | | | FKBP-like pe | nucleus | |
| AT1G62390 | 2.558 | 1.478 | | | Octicosapept | nucleus | |
| AT1G56110 | 2.487 | 2.054 | | | homolog of r | nucleus | |
| AT3G57150 | 2.457 | 2.000 | | | homologue o | nucleus | |
| AT5G50310 | 2.395 | 1.194 | | | Galactose ox | nucleus | |
| AT2G28490 | 2.324 | 1.127 | | | RmlC-like cu | extracellular | |
| AT5G13420 | 2.269 | 1.127 | | | Aldolase-typ | plastid | |
| AT4G10480 | 2.246 | 2.134 | | | Nascent poly | cytosol | |
| AT5G47210 | 2.182 | 1.054 | | | Hyaluronan / | nucleus | |
| AT4G39520 | 2.067 | 1.720 | | | GTP-binding | cytosol | |
| AT3G05060 | 2.006 | 2.211 | | | NOP56-like p | nucleus | |
| AT5G60980 | 1.936 | 1.478 | | | Nuclear trans | nucleus | |
| AT1G03880 | 1.846 | 1.677 | | | cruciferin 2, | endoplasmic reticulum | |

**Table S5**: Comparison of mitochondrial proteome in fmt and Col0 seedlings. With a Fold-Change cut-off above 1.5 and an adjusted p-value below 0.1, 72 proteins were found affected in fmt compared to Col0

| in *fmt* seedlings | Accession | Uniprot Name | Description | functional group | LogFC seedlings: FMT vs Col0 | adjp seedlings : FMT vs Col0 | spectral count / 100 aa in Col0 seedlings | LogFC infloresc : FMT vs Col0 | adjp infloresc : FMT vs Col0 |
|---|---|---|---|---|---|---|---|---|---|
| down | AT4G35490.1 | Q9SVW7 | mitochondrial ribo | mitorib | -1.854 | 0.072 | 3.441 | -0.217 | 1.000 |
| down | AT1G19520.1 | NFD5 | pentatricopeptide | mitorib | -1.472 | 0.001 | 2.943 | -0.313 | 0.772 |
| down | AT4G39880.1 | Q9SMR5 | Ribosomal protein | mitorib | -2.367 | 0.005 | 4.307 | 1.705 | 0.650 |
| down | AT1G07830.1 | Q94JQ7 | ribosomal protein | mitorib | -1.613 | 0.046 | 5.324 | 0.098 | 1.000 |
| down | AT2G20060.1 | Q8VY61 | Ribosomal protein | mitorib | -1.872 | 0.002 | 4.333 | -0.051 | 1.000 |
| down | AT5G44710.1 | Q8GXH5 | unknown protein | mitorib | -2.549 | 0.029 | 4.575 | 1.080 | 0.772 |
| down | **AT5G08530.1** | CI-51, NDUV1 | 51 kDa subunit of | OXPHOS-complex I | -0.658 | 0.001 | 15.844 | -0.471 | 0.403 |
| down | AT5G63510.2 | | gamma carbonic a | OXPHOS-complex I | -0.618 | 0.034 | 16.667 | -0.351 | 0.772 |
| down | AT3G48680.1 | GCAL2 | gamma carbonic a | OXPHOS-complex I | -0.783 | 0.011 | 14.453 | -0.372 | 0.772 |
| down | AT2G33220.1 | NDADB | GRIM-19 protein | OXPHOS-complex I | -1.044 | 0.009 | 16.550 | -0.052 | 1.000 |
| down | AT1G04630.1 | NDADA | GRIM-19 protein, | OXPHOS-complex I | -1.313 | 0.002 | 15.152 | -0.122 | 1.000 |
| down | AT3G12260.1 | NDUA6 | LYR family of Fe/S | OXPHOS-complex I | -0.590 | 0.095 | 24.311 | -0.355 | 0.772 |
| down | AT1G14450.1 | NDB3B | NADH dehydrogen | OXPHOS-complex I | -0.588 | 0.096 | 45.662 | -0.196 | 1.000 |
| down | ATMG00510.1 | NDUS2 | NADH dehydrogen | OXPHOS-complex I | -0.803 | 0.023 | 7.530 | -0.264 | 0.803 |
| down | ATMG00070.1 | NDUS3 | NADH dehydrogen | OXPHOS-complex I | -0.991 | 0.005 | 15.263 | -0.212 | 0.946 |
| down | AT3G62790.1 | NDS5B | NADH-ubiquinone | OXPHOS-complex I | -1.550 | 0.066 | 8.835 | -0.172 | 1.000 |
| down | AT5G67590.1 | NDUS4 | NADH-ubiquinone | OXPHOS-complex I | -1.112 | 0.011 | 13.203 | -0.603 | 0.650 |
| down | AT1G76200.1 | NDUB2 | unknown protein | OXPHOS-complex I | -5.332 | 0.001 | 7.246 | 0.000 | 1.000 |
| down | AT4G20150.1 | Q94AL6 | unknown protein | OXPHOS-complex I | -3.237 | 0.014 | 5.350 | -0.752 | 0.747 |
| down | AT1G67350.1 | Q9FYF8 | unknown protein | OXPHOS-complex I | -1.166 | 0.008 | 20.748 | -0.324 | 0.809 |
| down | AT3G27240.1 | CYC1A | Cytochrome C1 far | OXPHOS-complex III | -0.647 | 0.003 | 21.716 | 0.078 | 1.000 |
| down | **AT5G13430.1** | CIII RISP,UCRI1 | Ubiquinol-cytochro | OXPHOS-complex III | -1.061 | 0.001 | 12.255 | -0.023 | 1.000 |
| down | AT5G08670.1 | ATPBM | ATP synthase alph | OXPHOS-complex V | -0.650 | 0.000 | 72.842 | -0.228 | 0.401 |
| down | ATMG00410.1 | ATP61 | ATPase, F0 comple | OXPHOS-complex V | -3.237 | 0.014 | 1.126 | 0.000 | 1.000 |
| down | AT5G13450.1 | ATPO | delta subunit of M | OXPHOS-complex V | -0.636 | 0.013 | 21.709 | -0.179 | 0.876 |
| down | AT2G33040.1 | ATPG3 | gamma subunit of | OXPHOS-complex V | -0.669 | 0.004 | 18.462 | -0.095 | 1.000 |
| down | ATMG00640.1 | MI25 | hydrogen ion trans | OXPHOS-complex V | -0.774 | 0.029 | 15.451 | 0.240 | 1.000 |
| down | ATMG00480.1 | YMF19 | Plant mitochondria | OXPHOS-complex V | -0.725 | 0.031 | 20.675 | -0.052 | 1.000 |
| down | AT2G44350.2 | CISY4 | Citrate synthase fa | TCA | -0.604 | 0.001 | 22.128 | -0.035 | 1.000 |
| down | AT4G35260.1 | IDH1 | isocitrate dehydro | TCA | -0.656 | 0.001 | 19.619 | 0.097 | 1.000 |
| down | AT2G17130.1 | IDH2 | isocitrate dehydro | TCA | -0.727 | 0.003 | 14.532 | -0.421 | 0.758 |
| down | AT3G07480.1 | Q9SRR8 | 2Fe-2S ferredoxin-like superfamily... | | -0.915 | 0.038 | 12.788 | -0.026 | 1.000 |
| down | AT4G01660.1 | ABC1 | ABC transporter 1, Symbols: ATABC1... | | -1.739 | 0.003 | 2.087 | -0.117 | 1.000 |
| down | AT4G30490.1 | Q8L517 | AFG1-like ATPase family protein | | -2.646 | 0.019 | 1.006 | -0.443 | 1.000 |
| down | AT4G34310.1 | Q9SYZ6 | alpha/beta-Hydrolases superfamily ... | | -3.438 | 0.005 | 0.407 | 0.000 | 1.000 |
| down | AT2G25140.1 | CLPB4 | casein lytic proteinase B4, Symbol... | | -1.047 | 0.072 | 1.349 | -0.829 | 0.772 |
| down | AT5G10860.1 | CBSX3 | Cystathionine beta-synthase (CBS) ... | | -1.096 | 0.000 | 33.819 | -0.033 | 1.000 |
| down | AT5G12290.1 | DGS1 | dgd1 suppressor 1, Symbols: DGS1 | | -1.904 | 0.002 | 1.993 | -0.563 | 0.876 |
| down | AT3G24200.2 | Q2V3S9 | FAD/NAD(P)-binding oxidoreductase ... | | -1.565 | 0.001 | 3.696 | -0.212 | 1.000 |
| down | AT2G29080.1 | FTSH3 | FTSH protease 3, Symbols: ftsh3 | | -1.133 | 0.004 | 2.925 | -0.427 | 0.700 |
| down | AT5G07440.1 | DHE2 | glutamate dehydrogenase 2, Symbols... | | -1.153 | 0.000 | 32.036 | -0.073 | 1.000 |
| down | AT3G03910.1 | DHE3 | glutamate dehydrogenase 3, Symbols... | | -1.386 | 0.000 | 11.922 | -0.080 | 1.000 |
| down | AT5G56730.1 | PQQL | Insulinase (Peptidase family M16) ... | | -3.237 | 0.014 | 0.453 | -1.148 | 0.772 |
| down | AT1G79230.1 | STR1 | mercaptopyruvate sulfurtransferase... | | -0.816 | 0.005 | 10.466 | -0.255 | 0.772 |
| down | AT4G05020.2 | NDB2 | NAD(P)H dehydrogenase B2, Symbols:... | | -0.717 | 0.025 | 6.071 | 0.055 | 1.000 |
| down | AT1G28510.1 | Q9SGP9 | Optic atrophy 3 protein (OPA3) | | -1.412 | 0.034 | 6.043 | 0.071 | 1.000 |
| down | AT4G35850.1 | PP351 | Pentatricopeptide repeat (PPR) sup... | | -1.667 | 0.000 | 5.255 | -2.204 | 0.000 |
| down | AT5G14580.1 | PNP2 | polyribonucleotide nucleotidyltran... | | -1.267 | 0.048 | 1.110 | 0.437 | 0.807 |
| down | AT3G08860.1 | AGT23 | PYRIMIDINE 4, Symbols: PYD4 | | -1.645 | 0.018 | 2.079 | 0.000 | 1.000 |
| down | AT1G24610.1 | Q9FYK3 | Rubisco methyltransferase family p... | | -3.529 | 0.003 | 1.120 | -1.119 | 0.700 |
| down | AT3G15640.1 | CX5B1 | Rubredoxin-like superfamily protein | | -1.222 | 0.005 | 11.553 | 0.067 | 1.000 |
| down | AT2G04940.1 | Q9SI32 | scramblase-related | | -2.737 | 0.013 | 1.361 | -0.350 | 0.958 |
| down | AT5G37590.1 | | Tetratricopeptide repeat (TPR)-lik... | | -4.765 | 0.007 | 0.605 | -0.047 | 1.000 |
| down | AT1G15480.1 | PPR44 | Tetratricopeptide repeat (TPR)-lik... | | -1.787 | 0.018 | 1.459 | -0.485 | 0.918 |
| down | AT1G26460.1 | PPR58 | Tetratricopeptide repeat (TPR)-lik... | | -1.414 | 0.001 | 3.545 | -3.220 | 0.000 |
| down | AT3G58840.1 | PMD1 | Tropomyosin-related | | -1.294 | 0.085 | 2.725 | -0.178 | 1.000 |
| down | AT5G49210.1 | Q945P2 | unknown protein | | -4.765 | 0.007 | 1.709 | -0.443 | 1.000 |
| down | AT1G16000.1 | P93048 | unknown protein | | -3.052 | 0.002 | 7.752 | -1.894 | 0.772 |
| down | AT2G20390.2 | Q9SK63 | unknown protein | | -3.003 | 0.032 | 2.004 | 2.641 | 0.700 |
| down | AT5G23200.1 | Q9FMY0 | unknown protein | | -1.379 | 0.031 | 2.757 | 0.248 | 1.000 |
| down | AT1G72020.1 | Q9C7G5 | | | -2.041 | 0.072 | 4.811 | 2.641 | 0.700 |
| up | AT3G22370.1 | AOX1A | alternative oxidase | AOX | 3.453 | 0.000 | 0.565 | 2.647 | 0.277 |
| up | AT3G22360.1 | AOX1B | alternative oxidase | AOX | 3.772 | 0.000 | 0.205 | 2.726 | 0.626 |
| up | AT3G27620.1 | AOX1C | alternative oxidase | AOX | 3.466 | 0.000 | 0.203 | 2.170 | 0.687 |
| up | AT3G47930.1 | GLDH | L-galactono-1,4-la | OXPHOS-complex I | 1.008 | 0.000 | 10.219 | 0.855 | 0.000 |
| up | AT4G08900.1 | ARGI1 | arginase | | 0.869 | 0.003 | 6.238 | 0.047 | 1.000 |
| up | AT4G08870.1 | ARGI2 | Arginase/deacetylase superfamily p... | | 0.886 | 0.000 | 13.953 | -0.108 | 1.000 |
| up | AT5G55200.1 | MGE1 | Co-chaperone GrpE family protein | | 1.317 | 0.002 | 2.870 | 0.579 | 0.700 |
| up | AT4G17650.1 | Q8LAA4 | Polyketide cyclase / dehydrase and... | | 1.593 | 0.037 | 1.042 | 0.624 | 1.000 |
| up | AT3G22330.1 | RH53 | putative mitochondrial RNA helicas... | | 0.958 | 0.009 | 2.273 | 0.262 | 0.772 |
| up | AT5G54100.1 | Q9LVW0 | SPFH/Band 7/PHB domain-containing ... | | 0.860 | 0.028 | 3.408 | 1.285 | 0.001 |
| up | AT4G00026.1 | Q2V3L9 | unknown protein | | 1.973 | 0.025 | 0.620 | -0.049 | 1.000 |

**Table S6**: DNA constructs and plamids

| Construct | Plasmid | Sequence origin | Vector | reference |
|---|---|---|---|---|
| FMT-GFP | pAM557 | cDNA A. thaliana | pB7FWG2 | this work |
| GFP-FMT | pAM549 | cDNA A. thaliana | pK7WGF2 | this work |
| MS2-GFP | pAM495 | Michaud_PNAS2014 | pB7FWG2 | this work |
| beta11 | pAM570 | Addgene #97398 | pMDC232 (into AscI) | this work |
| beta11-FMT | pAM574 | cDNA A. thaliana | pAM570 | this work |
| beta11-GAPDH | pAM588 | Michaud_Bioch2014 | pAM570 | this work |
| beta11-TOM20 | pAM576 | cDNA A. thaliana | pAM570 | this work |
| beta11-UGPase | pAM616 | cDNA A. thaliana | pAM570 | this work |
| GFP$_{1_10}$-RPL18 | pAM602 | Addgene #97390 + cDNA A. thaliana | pMDC32 | this work |
| GFP$_{1_10}$-TOM5 | pAM573 | Addgene #97390 + cDNA S. cerevisae | pMDC32 | this work |

Michaud_PNAS2014: Michaud M., Ubrig E., Filleur S., Erhardt M., Ephritikhine G., Marechal-Drouard L. and Duchene A.M. (2014) Differential targeting of VDAC3 mRNA isoforms influences mitochondria morphology. PNAS 111: 8991-8996

Michaud_Bioch2014: Michaud M., Marechal-Drouard L. and Duchene A.M. (2014) Targeting of cytosolic mRNA to mitochondria: Naked RNA can bind to the mitochondrial surface. Biochimie 100: 159-166

**Table S7**: oligonucleotides used in qPCR and RT-qPCR

| Name | TAIR AGI | gene location | protein location | Primer (5' -> 3') | Primer 2 (5' -> 3') | Figure |
|---|---|---|---|---|---|---|
| NDHH | ATCG01110 | chloro | chloro | TCCGGATAAACCCCAATTTA | AGAACGGGTTGAAGGAGTTG | 7A |
| 16S | ATCG01210; ATCG00920 | chloro | chloro | GATCGGAAAGAACACCAACG | CCCCTAGCTTTCGTCTCTCA | 7A |
| 18S | ATMG01390 | mito | mito | CCTTGAGCTAGGAGCCTCTTT | CATGCAAGTCGAACGTTGTT | 7A |
| COX2 | ATMG00160 | mito | mito | AATAAACGTGATTGACCCAATTCT | TCCGATGAGCAGTCACTCAC | 7A |
| ACTIN1 | AT2G37620 | nuclear | cyto | CATCTTGGCCTCCCTCAGTA | GAGTAAACAAGTGATGGGACTGTG | 7A |
| 18S | 18S RRNA | nuclear | cyto | AAACGGCTACCACATCCAAG | ACTCGAAAGAGCCCGGTATT | 7A |
| 18S | ATMG01390 | mito | mito | CATGCAAGTCGAACGTTGTT | CCTTGAGCTAGGAGCCTCTTT | 7B |
| COX1 | ATMG01360 | mito | mito | GTAGCTGCGGTGAAGTAGGC | CTGCCTGGATTCGGTATCAT | 7B |
| COX2 | ATMG00160 | mito | mito | TCCGATGAGCAGTCACTCAC | AATAAACGTGATTGACCCAATTCT | 7B |
| NAD4 | ATMG00580 | mito | mito | AATACCCATGTTTCCCGAAG | TGCTACCTCCAATTCCCTGT | 7B |
| COB | ATMG00220 | mito | mito | TGCCGGAATGGTATTTCCTA | GCCAAAAGCAACCAAAACAT | 7B |
| RPL2 | ATMG00560 | mito | mito | CCGAAGACGGATCAAGGTAA | CGCAATTCATCACCATTTTG | 7B |
| RPS3 | ATMG00090 | mito | mito | CCGATTTCGGTAAGACTTGG | AGCCGAAGGTGAGTCTCGTA | 7B |
| GAPDH cyto | AT1G13440 | nuclear | cyto | AGGCTGCTGCTCACTTGAA | AACATGGGCGCATCTTTG | 7B; 8 |
| RPL12 cyto | AT2G37190 | nuclear | cyto | GACGTGTACGTCCGAGTAACC | GACCGATTTTGGGAGCTAGA | 7B; 8 |
| VDAC3 CDS | AT5G15090 | nuclear | mito | CACTGAAATCGGCAAAAAGG | TGTTCCCGTTGTAGTGATCG | 8 |
| VDAC3 long | AT5G15090 | nuclear | mito | TCCATATCTTTTACTTGGTTCTCTCTT | GGGAACTCCAAATGGAACAA | 8 |
| CI-51 | AT5G08530 | nuclear | mito | GAACAGGTTGGCTTTGGATG | TGGTTACCTCCTGCAGCATA | 8 |
| CIII-RISP | AT5G13430 | nuclear | mito | CCTTGCCTAATGCTGGTGA | ATATCGTAATGTGATCCGTGACA | 8 |
| CI-7.5 | AT1G67785 | nuclear | mito | TCTTCATCAAGACGAGAAGCAA | CCGTAGCAGCTCTGCTCTAAC | 8 |
